# Obesity medication lorcaserin requires brainstem GLP-1 neurons to reduce food intake in mice

**DOI:** 10.1101/2022.05.06.490598

**Authors:** Stefan Wagner, Daniel I. Brierley, Alasdair Leeson-Payne, Wanqing Jiang, Raffaella Chianese, Brian Y. H. Lam, Georgina K. C. Dowsett, Claudia Cristiano, David Lyons, Frank Reimann, Fiona M. Gribble, Giles S.H. Yeo, Stefan Trapp, Lora K. Heisler

## Abstract

Overweight and obesity are rapidly becoming the “new normal” in developed countries, which promotes a widespread negative impact on human health. Amongst recently developed obesity medications are the serotonin 2C receptor (5-HT_2C_R) agonist lorcaserin and glucagon-like peptide-1 receptor (GLP-1R) agonists, but the brain circuits employed by these medications to produce their therapeutic effects remain to be fully defined. 5-HT_2C_Rs and GLP-1Rs are widely distributed in the brain, including in the key homeostatic region the nucleus of the solitary tract (NTS) where GLP-1 is produced by preproglucagon (PPG^NTS^) neurons. PPG^NTS^ cells were profiled using histochemistry and single nucleus RNA sequencing (Nuc-Seq) of mouse brainstem. Transcriptomic analyses revealed 5-HT_2C_R expression was widespread in PPG^NTS^ clusters. Demonstrating the functional significance of this co-expression, lorcaserin required PPG^NTS^ to reduce food intake. Analysis of second order neurons revealed that local GLP1-R neurons within the NTS are necessary for 5-HT_2C_R^NTS^ food intake suppression. In contrast, GLP-1R^NTS^ were not required for GLP-1R agonist liraglutide and exendin-4’s short term feeding reduction, suggesting scope for lorcaserin and GLP1-R agonist combination therapy. In support of this, lorcaserin+liraglutide and lorcaserin+exendin-4 produced greater reductions in food intake when administered in combination as compared to monotherapies. These data provide insight into the therapeutic mechanisms of lorcaserin and identify a combination strategy to improve the therapeutic profile of lorcaserin and GLP1-R agonists.

## Main Text

The prevalence of obesity has reached epidemic levels, with approximately 40% of the global adult population now estimated to be overweight or living with obesity. This has profound health implications because obesity is a major risk factor for multiple chronic diseases. Medications developed to treat obesity include agonists for G-protein coupled receptors serotonin 2C (5-HT_2C_R), glucagon-like peptide-1 (GLP-1R), and melanocortin4 (MC4R) which reduce bodyweight primarily by suppressing food intake. However, the mechanisms underlying the therapeutic food intake suppressive effects of these medications are still being defined. In particular, it is unclear whether the brain circuits targeted by these agonists are entirely separate entities, or whether their pathways converge to drive weight loss by potentiation of satiation and/or satiety. To foster precision medicine, a better understanding of the neurophysiological control of appetite is essential to more effectively treat individuals with obesity. Clarifying these circuits may also drive drug discovery efforts for more effective new medications or combinations of licensed medications.

Consistent with the brain being the master coordinator of energy homeostasis, recent research indicates that the sites of action for the food intake suppressive effects of the 5-HT_2C_R agonist lorcaserin, GLP-1R agonists, and MC4R agonists reside within the CNS (1-8). For example, activation of brain GLP-1Rs by centrally injected GLP-1 or GLP-1R agonists effectively reduce feeding. This is mirrored by the more recent finding that activation of preproglucagon (PPG; product of the *Gcg* gene) expressing neurons in the nucleus of the solitary tract (PPG^NTS^) decreases food intake (9-12). PPG^NTS^ neurons constitute the primary brain source of GLP-1 (11), however the conditions under which these cells are recruited and their afferent neurocircuitry are poorly understood. In mice, PPG^NTS^ neurons express leptin receptors, and also receive substantial serotonergic input (13-15). In rats, GLP-1 immunoreactive neurons express serotonergic 5-HT_2C_Rs, though this has not yet been demonstrated in mice (7). However, 5-HT receptor agonist treatment in mice illustrates that 5-HT’s effects on PPG^NTS^ neuron activity is both excitatory via the G_q_-coupled 5-HT_2_R family, and inhibitory via the G_i_-coupled 5-HT_1A_R subtype (15). Here we examine whether PPG^NTS^ neurons express 5-HT_2C_Rs, and the functional implications of this for therapeutic regulation of food intake in mice.

Previous reports indicate that food intake reduction by the 5-HT_2C_R agonist lorcaserin requires a functional brain melanocortin system. Specifically, it was shown that lorcaserin acts via arcuate nucleus of the hypothalamus (ARC) and NTS pro-opiomelanocortin (POMC) signalling to MC4Rs to decrease food intake (5, 6). POMC’s peptide products α-, β- and γ-melanocyte-stimulating hormone (MSH) are the endogenous agonists for MC4Rs, and activation of MC4Rs effectively decreases food intake (16, 17). Although MC4Rs are expressed in a variety of brain regions, MC4Rs localized within the paraventricular nucleus of the hypothalamus (PVH) have been extensively studied and found to modulate appetite and body weight (2-4). We previously reported that the expression of 5-HT_2C_Rs exclusively within the NTS and dorsal motor nucleus of the vagus (DMX) is sufficient for lorcaserin’s acute effect on food intake (5). Although POMC^NTS^ is required for lorcaserin to produce this acute effect on feeding, the majority of 5-HT_2C_R^NTS^ are not expressed in POMC neurons (5). Here we examine the hypothesis that PPG^NTS^ neurons express 5-HT_2C_R and are involved in the mechanism of lorcaserin-induced appetite suppression.

To investigate potential overlap between NTS 5-HT_2C_R, the central GLP-1 system, and MC4Rs, we performed single nucleus RNA sequencing (Nuc-Seq) from mouse brainstem (18) (Table 1), histochemical analyses, and pharmacological, genetic and chemogenetic manipulations. Nuc-Seq yielded 86 extracted glucagon (*Gcg*) gene expressing cells, of which 74 were neurons. These cells separated into three major clusters, two of which contained neurons with high and low expression of *Gcg*, respectively (Fig 1a-d). The high *Gcg* cluster (n=28; cluster 1) was excitatory (glutamatergic), which is consistent with previous research indicating overlap between PPG^NTS^ and glutamatergic neurons (19, 20). However, we also identified a cluster of neurons that expressed low levels of *Gcg* transcript (n=46; cluster 0) and this was of a primarily inhibitory phenotype (GABAergic/glycinergic). To examine whether 5-HT_2C_Rs are positioned to influence the activity of PPG^NTS^ neurons in mice, we evaluated receptor expression patterns in these neurons. *Htr2c* (5-HT_2C_R) expression was widespread in both clusters, with 67% of cluster 0 cells and 79% of cluster 1 cells expressing this receptor (Fig 1e). This high degree of overlap is consistent with a previous report in rats (7). In contrast, expression of *Mc4r* and *Mc3r* was very low in both clusters with only 3 cells expressing *Mc4r* and only 1 cell expressing *Mc3r* (Fig 1f,g). Furthermore, out of 86 *Gcg* expressing and 346 *Pomc* expressing cells, only 2 cells co-expressed both genes, demonstrating very limited overlap between these two cell populations. We next examined 5-HT_2C_R^NTS^ cells from Nuc-Seq (Table 1). In accordance with previous studies, we found widespread expression of 5-HT_2C_R, and a small proportion of these expressed POMC (5). These findings suggest that 5-HT_2C_Rs, but not MC4Rs or MC3Rs, are positioned to influence the activity of a large proportion of PPG cells.

**Table 1.**
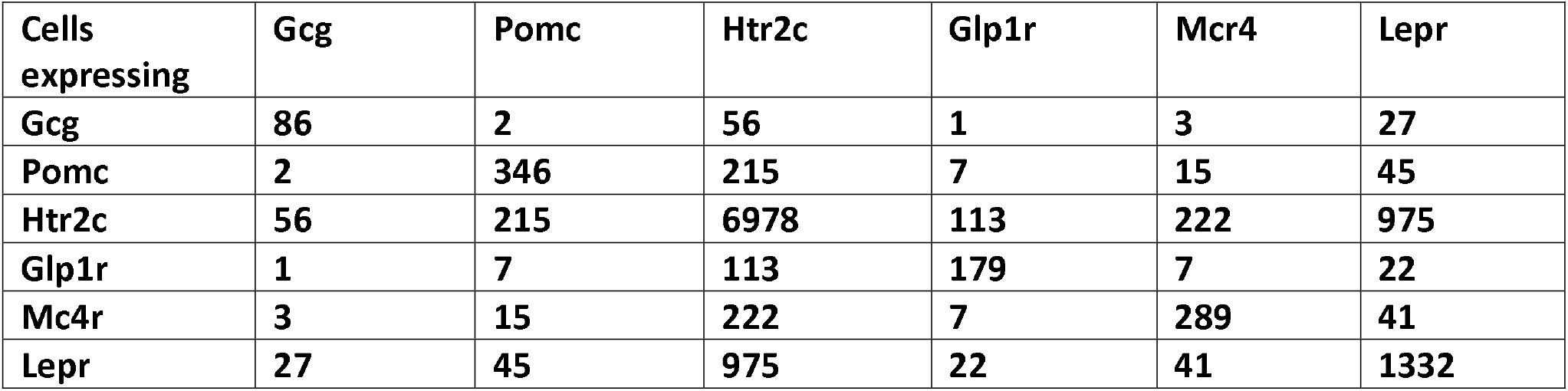

**Figure 1.**
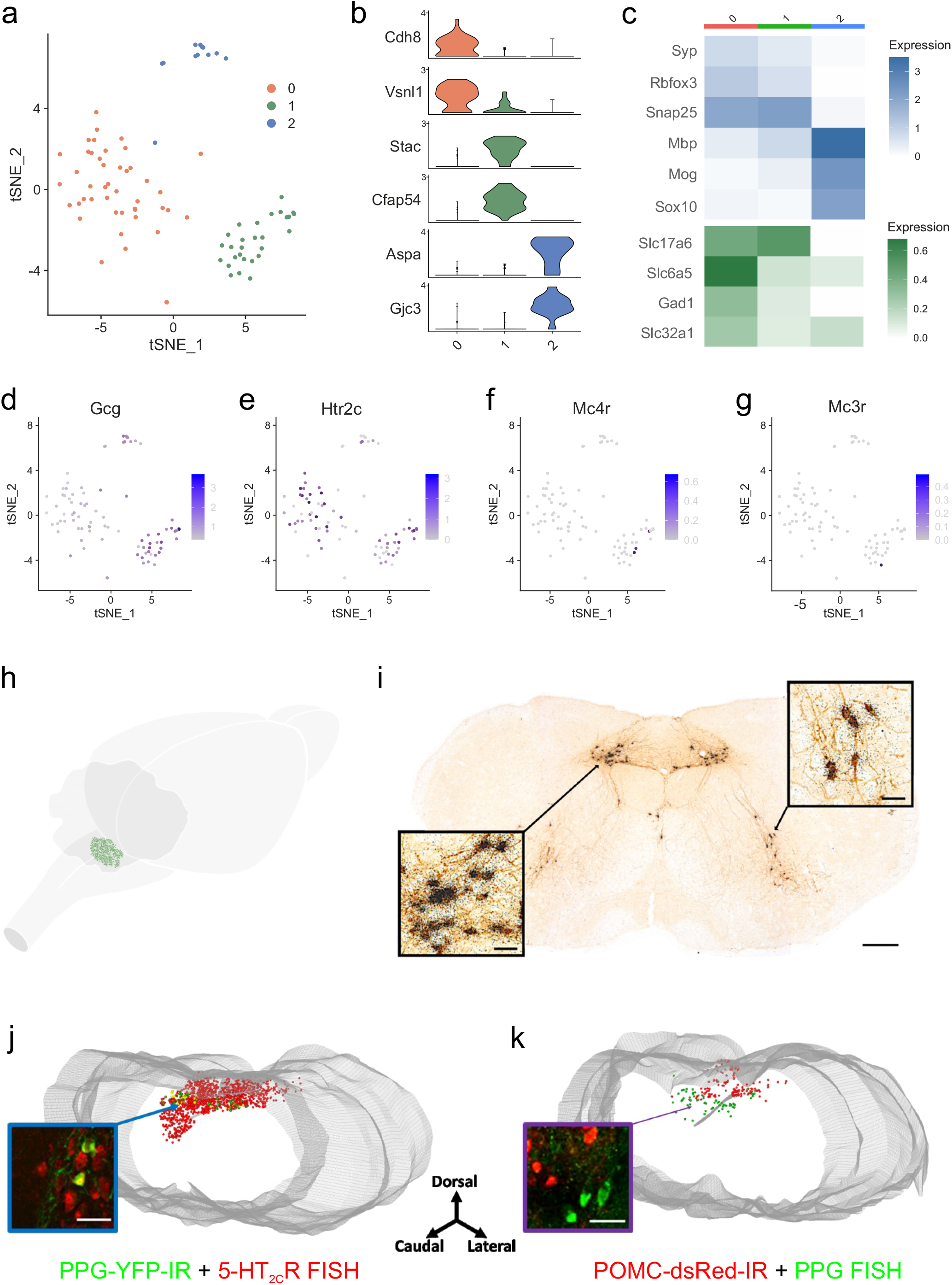
PPG ^NTS^ neurons co-express 5-HT_2C_Rs but not POMC or MC3/4Rs. (a) tSNE plot of 86 preproglucagon (PPG) cells (*Gcg*+) from the brainstem single nucleus RNA sequencing dataset grouped into three clusters. b) Violin plots showing the relative expression levels of two marker genes for each of the three PPG cell clusters. c) Heatmaps revealing the average relative expression of canonical neuronal and oligodendrocyte markers within each PPG cell cluster. (d-g) Feature plots showing the relative expression levels of *Gcg* (d) *Htr2c*(e), *Mc4r* (f) and *Mc3r* (g) in the PPG cells. (h) Three-dimensional (3D) schematic of the whole mouse brain illustrating the distribution of PPG neurons (green dots) within the nucleus of the solitary tract (NTS). (i) Representative photomicrographs of brainstem coronal section from a PPG mouse processed for immunohistochemistry for *GFP*-IR (brown stain) and in situ hybridization (ISH) with a ^35^S-labelled *Ppg*^*YFP*^ riboprobe (black grains) illustrating co-labelling within the NTS and intermediate reticular nucleus (IRt) (n=3 mice). (j) 3D rendering of the caudal to rostral NTS from PPG mice processed for IHC for *GFP*-IR (green cells) and 5-ht_2C_r mRNA with fluorescent ISH (FISH; red cells) illustrating single- and dual-labelling (yellow cells) (n=5 mice). (k) 3D rendering of the caudal to rostral NTS from *POMC*^*dsRED*^ mice processed for FISH for *Ppg*^*YFP*^ mRNA (green cells) and dsRed-IR (red cells) illustrating no overlap of POMC and PPG cells (n=4 mice). Grey tube is central canal (cc). Scale bars in i-k: 200 μm; insets 20 μm.

We next employed two transgenic mouse lines: mGlu-Venus (*Ppg* ^*YFP*^) and mGlu-Cre (*Ppg* ^*Cre*^) to further characterise PPG^NTS^ neurons and to confirm the Nuc-Seq data (11, 21). Anatomically, PPG neurons are expressed within the NTS and adjacent intermediate reticular nucleus (IRt) at the level of the area postrema and extend approximately 400 μm caudally (Figure 1h). Histochemical validation of the *Ppg*^*YFP*^ mouse line was performed using endogenous *Ppg* mRNA. Specifically, radioactive in situ hybridization histochemistry (ISHH) with a ^35^S *Ppg* riboprobe on NTS sections from *Ppg*YFP mice revealed that 97.9 ± 1.2% of *Ppg* mRNA positive cells exhibited YFP immunoreactivity (YFP-IR; using an anti-*GFP* antibody (21)) and 95.8 ± 1.7% of YFP-IR positive cells co-labelled with ^35^S *Ppg* (Figure 1i). In the IRt, 95.5 ± 1.7% of ^35^S *Ppg* positive cells exhibited YFP-IR and 92.9 ± 1.9% of YFP-IR positive cells co-labelled for ^35^S *Ppg* (Figure 1i). This near total overlap between YFP-IR and endogenous *Ppg* mRNA was replicated using fluorescent ISH (FISH) for *Ppg* (Supplementary Figure 1a-b). These data confirm that the *Ppg*^*YFP*^ mouse line and in situ methods for labelling *Ppg* mRNA may be used interchangeably to visualize PPG neurons.

We then used the *Ppg*^*YFP*^ line to confirm the Nuc-Seq findings. Using FISH for *Htr2c* co-localised with GPF-IR in *Ppg*^*YFP*^ mice revealed that 38.8 ± 2.8% of PPG cells were co-labelled for 5-HT2CR mRNA (Figure 1j). Though this rate of co-expression is lower than expected based on the Nuc-Seq method, these data corroborate the finding that 5-HT_2C_Rs are positioned to impact a substantial proportion of PPG^NTS^ neurons in mice. In agreement with the transcriptomics analysis (Table 1), 5-HT_2C_R-PPG neurons comprise a small proportion of the total 5-HT_2C_R^NTS^ population, as shown in a three-dimensional (3D) representation of the expression profile within the NTS (Figure 1j). To establish whether this anatomical localization identifies a previously uncharacterized population of 5-HT_2C_R^NTS^ cells, we examined co-expression of *Ppg* mRNA and POMC using dsRed in a *Pomc*^*dsRED*^ reporter line. We observed that PPG and POMC are separate populations of neurochemically-defined NTS cells (Figure 1k), corroborating a previous report utilizing ^35^S *Ppg* in *PomcGFP* mice (14) as well as the transcriptomics data presented here (Table 1). Taken together, these findings indicate that 5-HT_2C_Rs are positioned to directly influence the activity of approximately one-third to two-thirds of PPG^NTS^ neurons in mice.

We next investigated whether systemically administered lorcaserin activates PPG^NTS^ neurons. We first confirmed that 7.5 and 10 mg/kg i.p. lorcaserin significantly reduced chow intake at 1, 3 and 6 hours compared to vehicle treatment in wild type mice (Supplementary Figure 2a-c). We then determined that lorcaserin (10 mg/kg, i.p.) increased overall NTS neuronal activity compared to saline treatment as measured by FOS-IR (279 ± 58 cells vs 44 ± 16; Figure 2a). Furthermore, lorcaserin treatment induced significantly more FOS-IR specifically in YFP-IR PPG^NTS^ neurons relative to saline treatment (27.3 ± 6.3% vs 4.1 ± 2.5%; Figure 2b-h). These results demonstrate that a subset of PPG neurons, consistent with the proportion expressing 5-HT_2C_R as determined by FISH, is activated by lorcaserin. This finding extends our previous observation using *ex vivo* brainstem slices which demonstrated that approximately one third of PPG^NTS^ neurons show increased electrical activity upon 5-HT administration, and that this is mediated by 5-HT_2_ receptors (15).

**Figure 2.**
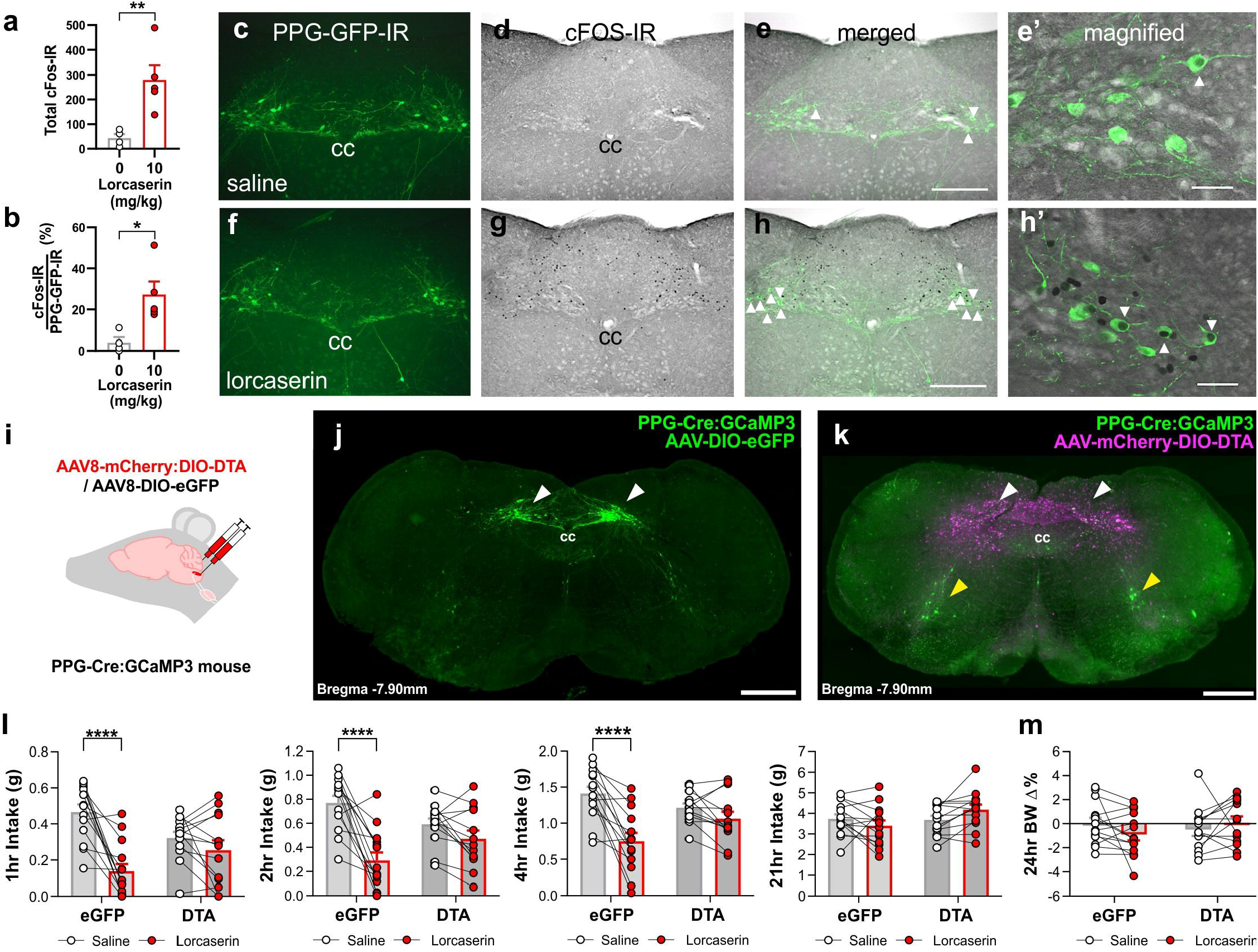
Lorcaserin requires PPG^NTS^ neurons to reduce food intake. (a-h) Lorcaserin-induced (10 mg/kg, i.p.) c-FOS immunoreactivity (IR) in PPG^NTS^ neurons in male and female *Ppg*^YFP^ mice (n = 10). (a) Lorcaserin increased the number of NTS c-FOS-IR cells (*t*(7) = 3.502, p = 0.010) and (b) the percentage of *GFP*-IR PPG^NTS^ neurons which were c-FOS-IR positive (Mann-Whitney U = 0, p=0.0159) compared to saline treated mice. (c-h) Photomicrographs of representative coronal NTS images of *GFP*-IR (green) and FOS-IR (black) positive cells following (c-e) saline or (f-h) lorcaserin treatment. (i) The nucleus of the solitary tract (NTS) of PPG-Cre:GCaMP3 mice was injected bilaterally with either AAV8-mCherry:DIO-DTA to ablate PPG neurons or with AAV8-DIO-e*GFP* as control. (j,k) representative photomicrographs showing the presence or absence of PPG^NTS^ neurons (white arrow heads), respectively, after transduction with control or DTA virus. Magenta shows spread of the DTA virus by visualising the Cre-independent transduction of mCherry. Yellow arrowheads indicate PPG^IRT^ neurons that are not reached by the virus. (l-n) Lorcaserin (7.5 mg/kg i.p.). significantly reduced cumulative dark cycle 1, 2, and 4 hour chow intake in male and female e*GFP*-transduced control (n=14) but not PPG^NTS^ DTA-ablated (n=14) *Ppg*^Cre^ mice (1hr: treatment x virus F (1,25) = 10.81, p=0.0030; 2hr: treatment x virus F_(1,25)_=11.93, p=0.0020; 4hr: treatment x virus F_(1,25)_= 11.35, p=0.0025). Lorcaserin did not significantly alter (o) 21 hour food intake or (p) 24 hour % change in body weight. cc, central canal. Scale bar in h 200 μm applies to c-h; scale bar in e’ and h’ 50 μm; scale bar in j,k 500 μm. Data are presented as mean ± SEM; * p < 0.05, ** p < 0.01, *** p < 0.001.

Having established that a subpopulation of PPG^NTS^ neurons both express 5-HT_2C_R and are activated by lorcaserin, we next examined whether these neurons are functionally necessary for the drug’s anorectic effect. A selective loss-of-function approach was employed to ablate PPG^NTS^ neurons in male and female *Ppg*^*Cre*^ mice by injecting either AAV8-mCherry-DIO-Diphtheria toxin subunit A (DTA) or control AAV8-DIO-enhanced green fluorescent protein (E*GFP*) into the NTS (11, 12)(Figure 2i-k; Supplementary Figure 2). Ablation of PPG^NTS^ neurons was verified in all mice by histological analysis (Supplementary Figure 2d-j). Chow intake was measured at 1, 2, 4 and 21 hours after dark onset in *ad libitum* fed mice. Lorcaserin induced a significant reduction in cumulative food intake during hours 1-4 in the control group, however this reduction was prevented by DTA ablation of PPG^NTS^ cells (Figure 2l-n). In patients, lorcaserin is taken twice daily, therefore as expected, one treatment with lorcaserin did not significantly impact 21 hour food intake or 24 hour body weight (Figure 2o,p). These findings identify PPG^NTS^ neuron activity as a necessary component of the food intake suppressive effect of lorcaserin in mice and suggest that pharmacological activation of approximately one-third of PPG neurons in the NTS is sufficient to elicit a significant reduction in food intake. Previous observations in rats using intracerebroventricular administration of the GLP-1 receptor antagonist exendin-9 showed a prevention of the anorexic effects of lorcaserin (7), consistent with our conclusion that lorcaserin activates PPG neurons in the brainstem to elicit hypophagia.

These experiments demonstrated that functional PPG neurons are necessary for the full hypophagic effect of lorcaserin. We next examined the mechanism of this effect, postulating that our anatomical data support a direct effect of 5-HT_2C_Rs on PPG^NTS^ neuron activity, promoting the local release of GLP-1. To test this hypothesis, we exclusively activated 5-HT_2C_R-expressing NTS neurons and established the necessity of GLP-1 release for their effects on food intake. Specifically, *5-HT2C*R^*Cre*^ mice received NTS injections of a viral cocktail comprising AAV8-DIO-hM3Dq-mCherry (or DIO-mCherry as a control) to evaluate the effect of 5-HT_2C_R^NTS^ neuronal activation, and the shRNA knockdown virus AAV1-shGlp1r-*GFP* (or AAV1-*GFP* as a control) to assess the importance of GLP-1R^NTS^ in feeding effects induced by chemogenetic activation of 5-HT_2C_R^NTS^ neurons (Figure 3). In situ hybridization histochemistry for *Glp1r* revealed that the majority of GLP-1Rs in the NTS (61.15 ± 11.27%), and a minority in the area postrema (15.54 ± 7.40%), were knocked down in mice injected with AAV1-shGlp1r compared with controls (Figure 3b-d, Supplementary Figure 3a). Mice were tracked for 2 months post-treatment. Similar to data in rats, knockdown of GLP-1R within the NTS had no effect on *ad libitum* food intake or body weight (Supplementary Figure 3b-e). These cohorts of mice were then injected with either clozapine-N-oxide (CNO) or saline, and food intake was measured up to 6 hours later (Figure 3e-h). 5-HT_2C_R^NTS^ neuron activation in mice expressing hMD3q (DREADD-Gq) showed a significant reduction in food intake 1-6 hours after administration of CNO compared to the control (Figure 3e-h). However, mice with GLP-1R^NTS^ knockdown did not show an anorectic response to CNO-induced DREADD-Gq activation of 5-HT_2C_R^NTS^ neurons (Figure 3e-h). These results confirm our previous findings that 5-HT_2C_R^NTS^ activation suppress food intake (5) and demonstrate that this response is dependent on the activation of GLP-1Rs expressed by NTS neurons. Considered with the results presented in Figure 2, the data from this GLP1-R knockdown experiments suggest that GLP-1R^NTS^ neurons are a target for 5-HT_2C_R-expressing PPG^NTS^ neurons.

**Figure 3.**
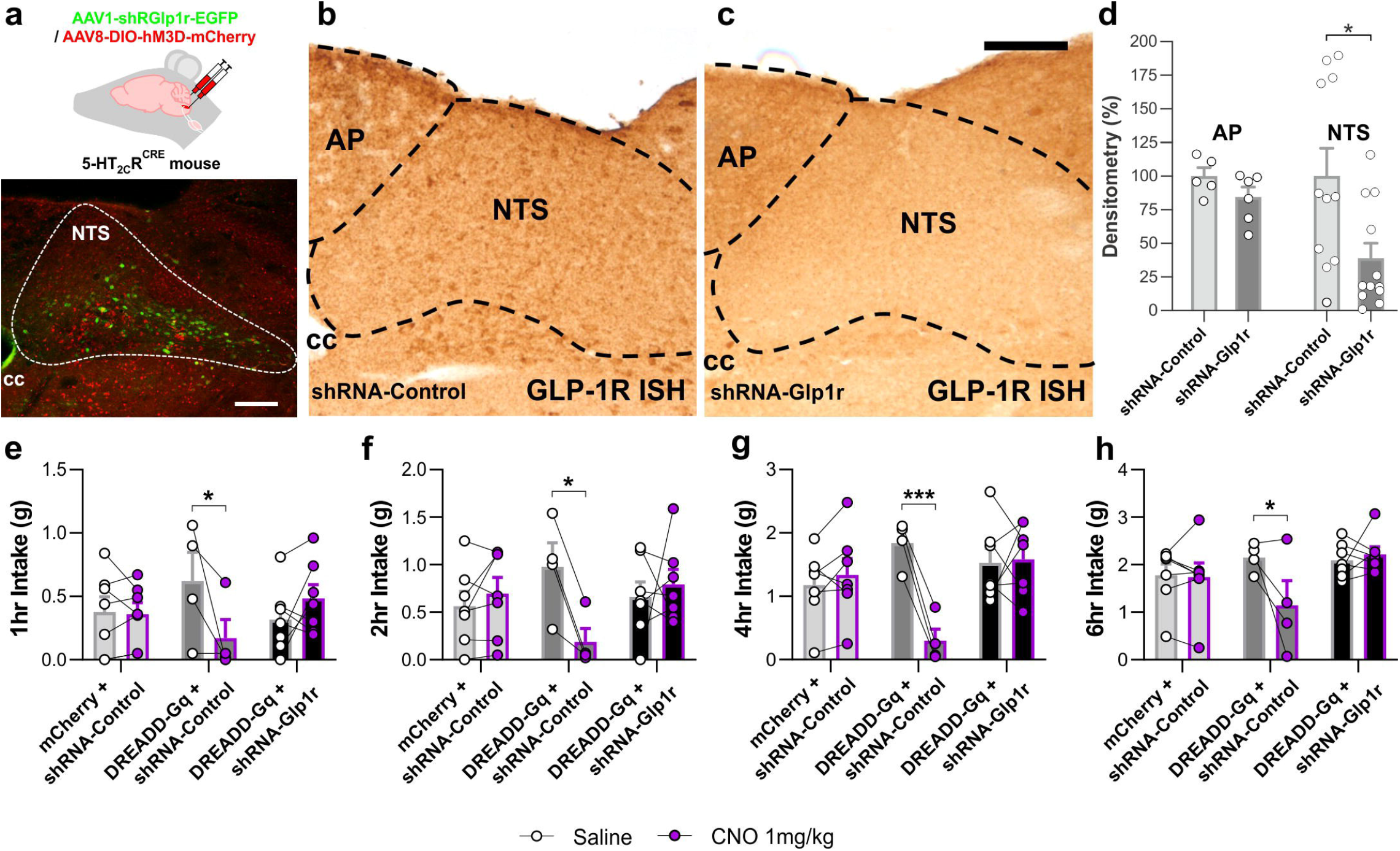
NTS^NTS^ GLP-1Rs are necessary for the anorectic effect of 5-HT_2C_R neuron activation. (a-h) The nucleus of the solitary tract (NTS) of 5-HT 2CR^CRE^ mice was injected with either DREADD-Gq (AAV8-hSyn-DIO-hM3Dq-mCherry) or mCherry control (AAV8-hSyn-DIO-mCherry) to allow targeted activation of 5-HT_2C_R cells, and simultaneously with shRNA-Glp1r (AAV1.U6.shRGlp1r07.CB7.E*GFP*.SV40) or control shRNA-Control (AAV1.CB7.Cl.e*GFP*.WPRE.rBG) to knock down GLP-1R expression (mCherry + shRNA-Control: n=7; DREADD-Gq + shRNA-Control: n=4; DREADD-Gq + shRNA-Glp1r: n=7). (a) Representative photomicrograph of injection site showing DREADD-Gq (red) and shRNA-Glp1r (green) in the NTS and schematic showing the injection site of viral delivery into the NTS. Scale bar 200μm. (b-d) Confirmation of Glp1r knockdown in NTS by *in situ* hybridisation in (b) shRNA-Control and (c) shRNA-Glp1r. Scale bar 50μm; CC, central canal. (d) quantification of knockdown by densitometry in area postrema (AP) and NTS. (e-h) These groups of mice were injected with CNO (1mg/kg) or saline and dark cycle food intake was measured at 1, 2, 4, and 6 hours following administration. Cumulative food intake was significantly decreased in DREADD-Gq + shRNA-Control mice but not DREADD-Gq + shRNA-Glp1r or mCherry + shRNA-Control mice at (e) 1 hour (AAV x treatment F_(2,15)_=5.533, p=0.0159), (f) 2 hours (AAV x treatment F_(2,15)_=4.949, p=0.0224), (g) 4 hours (AAV x treatment F_(2,15)_=13.44, p=0.0005), or (h) 6 hours (AAV x treatment F_(2,15)_=4.877, p=0.0234). Data presented as mean ± SEM; * p<0.05, ** p<0.01, *** p<0.001.

These results, combined with the earlier observation that lorcaserin requires POMC neurons in the arcuate nucleus signalling through MC4Rs for its intake suppressive effect (5, 6), suggests that lorcaserin exerts its anorectic effect by recruitment of either parallel or overlapping pathways. Consequently, we explored evidence for crosstalk between these pathways using neuroanatomical tracing and histochemistry to assess whether PPG^NTS^ neurons receive close appositions from 5-HT_2C_R^ARC^ cells, and whether PPG^NTS^ neurons express MC4Rs. To investigate this, a Cre-inducible AAV9-CAG-DIO-tdTomato tracer virus was injected into the ARC of *5-HT2CR*^*CRE:YFP*^ mice (Supplementary Figure 4a). We observed that 5-HT_2C_R^ARC^ axons robustly innervate the NTS (Supplementary Figure 4b). Using *Ppg* FISH, we found that 5-HT_2C_R^ARC^ axons were in close apposition to PPG^NTS^ neurons (Supplementary Figure 4b, inset). This pattern was not observed in *5-HT2CR*^*CRE:YFP*^ mice injected with AAV9-CAG-DIO-tdTomato into another hypothalamic sub-region, the posterior hypothalamus (Supplementary Figure 4c,d). Within the ARC, the subset of 5-HT_2C_Rs that are both necessary and sufficient to modulate the anorectic effect of 5-HT_2C_R agonists are expressed by POMC neurons (22, 23). POMC peptides signal through the MC4R to impact feeding (24, 25). Using a *Mc4r* null line, we confirmed that lorcaserin requires functional MC4Rs to reduce food intake (Figure 4a). To further examine whether PPG^NTS^ neurons express MC4Rs, we performed *Ppg* FISH in *Mc4r GFP* mice. We observed that a small proportion of PPG cells (13.5±1.3%) co-localise with *GFP*-IR from this MC4R reporter (Supplementary Figure 4e), consistent with our Nuc-Seq data demonstrating *Mc4r* expression in only three of the 86 identified PPG cells (Figure 1f). Our anatomical and transcriptomic data therefore suggest that lorcaserin may activate subsets of PPG^NTS^ neurons directly (via PPG^NTS^-5-HT_2C_Rs) and indirectly (via 5-HT_2C_R^ARC^ to PPG^NTS^), but that melanocortin signalling via MC4Rs expressed on PPG neurons may only be a minor contributor to this effect.

**Figure 4.**
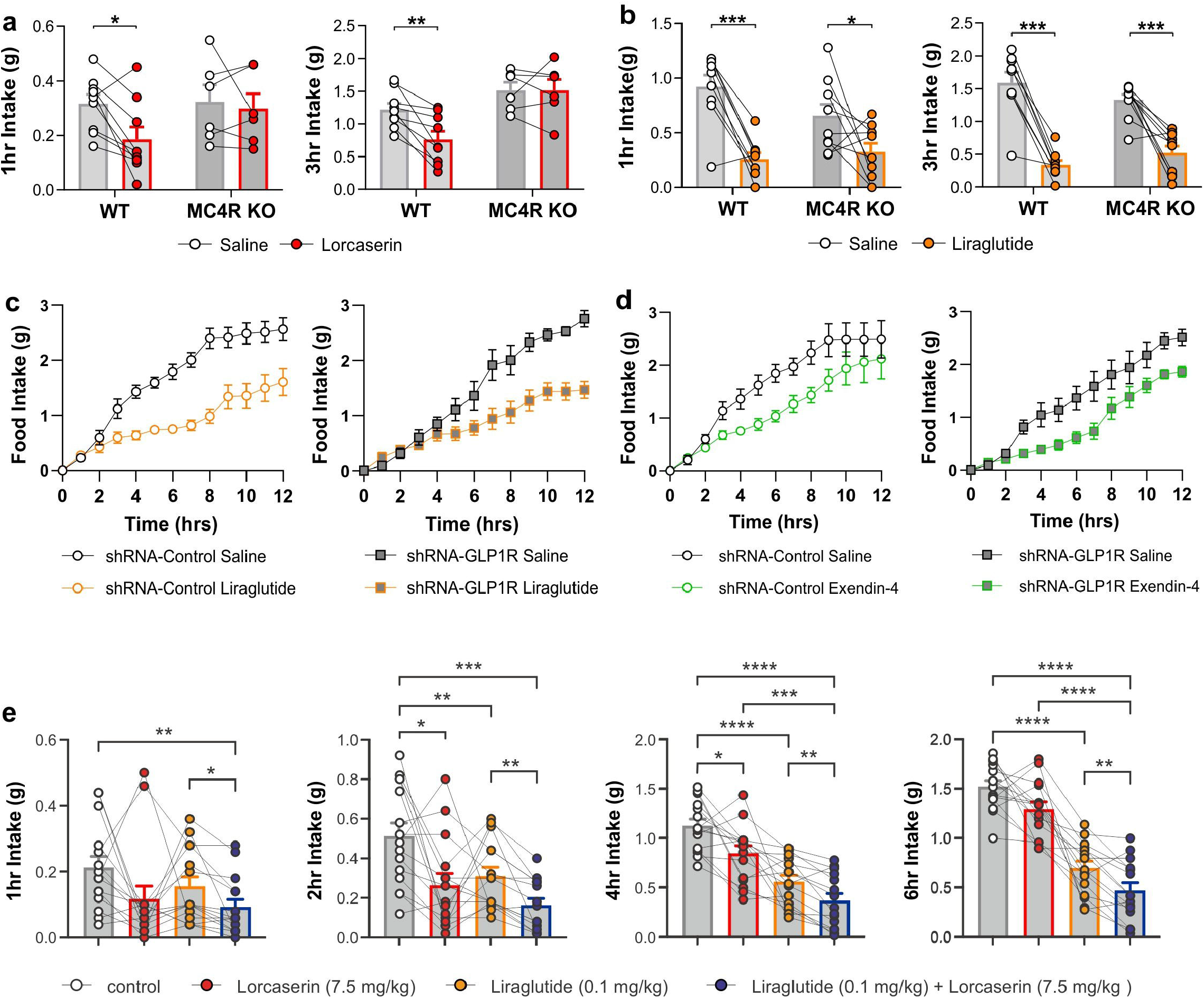
Combination of lorcaserin and GLP-1R agonist produces augmented reductions in food intake. (a) Lorcaserin (7.5 mg/kg, i.p.) signficantly reduced (a) 1 and 3 hour cumulative dark cycle food intake in male and female wild type (n=9), but not *Mc4r* knockout (n=7) mice. (1hr: treatment x genotype F_(1,13)_=2.464, p=0.1405; 3hr: treatment x genotype F_(1,13)_=6.898, p=0.0209). (b) In contrast, liraglutide (0.1 mg/kg, s.c.) significantly reduced 1 and 3 hour cumulative food intake in both adult wild type (n=8) and MC4R null (n=8) mice (1hr: treatment x genotype F_(1, 17)_ = 2.932, p=0.1050; 3hr: treatment x genotype F_(1, 17)_ = 4.062, p=0.0600). (c-d) Liraglutide (0.1 mg/kg, s.c.) significantly reduced cumulative food intake in both shRNA-Control (treatment F_(1,15)_=36.07, p=0.0001) and shRNA-Glp1r NTS knockdown mice (treatment F_(1,12)_=11.26, p=0.006) over 12 hours. Likewise, exendin-4 (0.008 mg/kg, i.p.) significantly reduced cumulative food intake in both shRNA-Control (treatment F_(1,16)_=5.52, p=0.03) and shRNA-Glp1r NTS knockdown mice (treatment F_(1,12)_=7.79, p=0.02) over 12 hours. (e) Individually effective anorectic doses of lorcaserin (7.5 mg/kg, i.p.) and liraglutide (0.1mg/kg) were tested for their ability to additively suppress food intake over 1-6 hours in male and female wild type (n=15). Combination treatment produced a greater reduction in food intake compared to lorcaserin and/or liraglutide alone at all timepoints (1hr: treatment F_(1.7,23.2)_=3.943, p=0.0404; 2hr: treatment F_(1.8, 25.4)_=11.41, p=0.0004; 4hr: treatment F_(2.2, 30.6)_=37.33, p<0.0001; 6hr: treatment F_(2.2, 31.4)_=73.72, p<0.0001). Data presented as mean ± SEM; * p<0.05, ** p<0.01, *** p<0.001, **** p<0.0001.

We tested this hypothesis by assessing the effect of the α-MSH analogue / MC4R agonist melanotan-II (MTII) on food intake and body weight in *PPG*^*Cre*^ mice with ablated (AAV8-mCherry-DIO-DTA) or functional (AAV8-DIO-E*GFP*) PPG^NTS^ cells. Using the same conditions employed in the lorcaserin treatment experiments, we observed that MTII (3 mg/kg, i.p.) induced a robust anorectic effect over 1 to 4 hours in both control e*GFP* and PPG^NTS^ ablated mice (Supplementary Figure 4f-i). The apparently more modest effect of MT-II on 21hr intake and 24hr bodyweight in PPG^NTS^ ablated mice (Supplementary Figure 4j-k) is most likely a non-specific interaction, driven by a larger post-treatment refeeding response in this group, similar to that seen in this ablation model following a prolonged fast (11, 12). Our finding that PPG^NTS^ neurons are not necessary for the acute anorectic effect of MTII is in line with the small proportion of anatomical overlap and limited expression of MC4R expression on PPG^NTS^ neurons, and with a previous *ex vivo* study employing patch clamp electrophysiology that failed to detect effects of MTII on the electrical activity of individual PPG neurons (26). Overall, these findings argue against any substantial direct effect of MC4R activation on PPG^NTS^ neurons, and are consistent with earlier reports that identified the population of MC4Rs that are sufficient to promote MTII’s effects on 3 hour food intake as residing within the PVH (3, 4). Our prior and current findings thus demonstrate that lorcaserin reduces food intake by activating both POMC→MC4R neuron driven hypophagia (5, 6) and PPG→GLP-1R neuron driven hypophagia, and that the loss of either pathway is sufficient to negate the food intake suppressive effects of lorcaserin.

We next investigated whether systemically-administered GLP-1R agonists require MC4Rs and/or GLP-1R^NTS^. We first performed a dose-response study to establish effective doses of the GLP-1R agonists liraglutide and exendin-4 in the reduction of food intake (Supplementary Figure 5a-f), then assessed the intake suppressing effects of these doses in MC4R knockout and GLP-1R knockdown mouse models. In contrast to lorcaserin, GLP-1R agonists liraglutide and exendin-4 did not require functional MC4Rs to impact feeding in mice (Figure 4a-b and Supplementary Figure 5g-h), which corroborates an earlier report in humans with MC4R mutations who were prescribed liraglutide (27). We then examined the response of mice lacking GLP-1R^NTS^ to systemically administered GLP-1R agonists liraglutide and exendin-4. In contrast to the loss of effect of chemogenetic activation of 5-HT_2C_R^NTS^ neurons in mice following knockdown of GLP-1R^NTS^, both liraglutide and exendin-4 largely retained their acute hypophagic effects in this mouse model (Figure 4c-d). This suggests that in mice, NTS GLP-1Rs are primarily a target for GLP-1 released from brain PPG neurons, rather than for systemically administered GLP-1R agonists. These findings are in line with a previous report in rat that GLP-1R expressing neurons in the area postrema, but not in the caudal NTS, display cFos immunoreactivity upon systemic administration of liraglutide (28). Similarly, the acute anorectic effect of liraglutide is maintained in rats with GLP1-R^NTS^ knockdown, though GLP1-R^NTS^ are required for the prolonged effect of liraglutide (29). Another possible explanation for GLP1-R^NTS^ knockdown and MC4R null mice largely responding to the anorectic effect of liraglutide and exendin-4 is that GLP-1R agonism is reported to inhibit GI motility (30). Thus, we conclude that both PPG^NTS^→GLP1R^NTS^ and POMC^NTS/ARC^→MC4R neurons are independent targets for the anorectic effects of lorcaserin.

Based on our results suggesting the acute hypophagic pathways recruited by lorcaserin and those recruited by systemic GLP-1R agonists are largely separate, we hypothesized that these compounds could be suitable for combination therapy. Therefore, in a final proof-of-concept experiment, we tested whether combining lorcaserin with liraglutide or exendin-4 could elicit greater reductions in food intake as compared to when these drugs were administered alone. Supporting our hypothesis, combined administration of individually effective doses of lorcaserin and liraglutide resulted in a significantly greater anorectic effect compared with administration of liraglutide alone at all time points tested (Figure 4e). To investigate whether this effect may be additive or synergistic, we next tested doses of lorcaserin and exendin-4 that do not significantly alter feeding when administered as monotherapies. While these doses were ineffective alone, they produced a robust reduction in food intake when administered in combination (Supplementary Figure 5i-k). These findings provide compelling support for a dual GLP-1R and 5-HT_2C_R agonist strategy for obesity treatment.

5-HT_2C_R, GLP-1R and MC4R agonists have been developed for human obesity treatment. However, the mechanisms underlying the therapeutic effects of these medications are still being defined. Here we report that PPG^NTS^ neurons function as a necessary hub through which lorcaserin achieves its full therapeutic benefit. Defining the circuits through which obesity medications promote their therapeutic effects will provide crucial insights into the appropriate choice of medicines for individuals with obesity. For example, patients with genetic MC4R alterations are unlikely to respond well to lorcaserin or the clinical MC4R agonist setmelanotide, but may have a better therapeutic response to liraglutide or semaglutide. All three types of medication would be expected to produce a therapeutic effect in patients with common obesity without genetic alterations in these circuits. Importantly, we now provide a strong rationale for clinical investigation of the combination of GLP1R agonists with lorcaserin, which has the potential for greater therapeutic efficacy and/or tolerability profile than either monotherapy alone.

## Author contributions

The project was conceived by ST and LKH. Data generation was led by SW and DIB with the assistance of AL-P, WJ, RC, CC, and DL. FR and FMG provided breeding pairs from *Ppg*^*YFP*^ Glu^*CRE*^ mouse lines. GY, BL and GD produced single nucleus sequencing data. The manuscript was drafted by ST and LKH with input from all other authors.

## Acknowledgements

The authors thank Dr Alasdair S. Garfield and staff within the University of Aberdeen Medical Research Facility and the Microscopy Facility for their technical assistance. pAAV-mCherry-flex-dtA was a gift from Naoshige Uchida and AAV1.U6.shRGlp1r07.CB7.E*GFP*.SV40 was a gift from Matthew Hayes. Work was supported by the Biotechnology and Biological Sciences Research Council (BB/K001418/1, BB/NO17838/1 to LKH and BB/S017593/1 to BYHL), the European Foundation for the Study of Diabetes (Merck Sharpe Dohme grant to ST), the Medical Research Council (MC/PC/15077 to LKH, MR/N02589X/1 to ST and MRC_MC_UU_12012/3 to FR and FMG, and MC_UU_00014/1 to GSHY), the NIH (R01 DK095757 to ST) and the Wellcome Trust (WT081713, WT098012 to LKH, 106262/Z/14/Z, 106263/Z/14/Z to FR and FMG, and 223279/Z/21/Z to DIB). Next-generation sequencing was performed by IMS Genomics and transcriptomics core facility, which is supported by the MRC (MC_UU_00014/5) and the Wellcome Trust (208363/Z/17/Z), and the Cancer Research UK Cambridge Institute Genomics Core. GKCD is funded by a BBSRC CASE 4-year PhD studentship, co-funded by Novo Nordisk. WJ is supported by a UCL GRS scholarship and a CSC scholarship from the Chinese Government.

## METHODS

### Mice

Adult male and female *Ppg*^*YFP*^ (31), *Glu*^*CRE*^ (32), *5-HT2CR* ^*CRE*^, *5-HT2CR*^*CRE:YFP*^ (5, 33), *Pomc* ^*DsRED*^ (5, 34), MC4R^GFP^ (Jackson Labs, B6.Cg-Tg (*Mc4r GFP*-MAPT/Sapphire)21Rck/J) and loxtb MC4R null (3) mice were used. Mice were group-housed whenever possible and kept on a 12 hour light/12 hour dark cycle with *ad libitum* access to chow and water unless otherwise stated.

### Single nucleus RNA sequencing

The clustered mouse brainstem single nucleus RNA sequencing data (generation described in (18)) is available from NCBI Gene Expression Omnibus Series (GSE168737). Analysis was performed using the Seurat package version 4.0.3 (35, 36). To remove the nuclei with low quality reads, nuclei with less than 1000 reads or 200 features were removed. Additionally, nuclei with more than 6000 features, which may be doublets, were also removed. Nuclei expressing at least one 1 raw UMI count of *Gcg* gene were extracted for analysis. This subset was then log-normalised and scaled, followed by principal component analysis (PCA). It then underwent non-linear dimensional reduction using T-distributed Stochastic Neighbour Embedding (tSNE) and unsupervised clustering using Louvain algorithm. The violin plots, feature plots and heatmaps were generated using the Seurat package.

### Tissue processing

Tissue was processed as previously described (5, 37, 38). Briefly, under deep terminal anaesthesia, mice were transcardially perfused with phosphate buffered saline (PBS) followed by 10% neutral buffered formalin (Sigma-Aldrich). Brains were extracted, post-fixed in 10% neutral-buffered formalin at 4°C, cryoprotected in 30% sucrose in PBS at 4°C and then sectioned coronally on a freezing sliding microtome at 25 μm. A total of five equal series was collected for each brain. For subsequent immunohistochemistry and in situ hybridization/immunohistochemistry (below), a single series was processed resulting in 125 μm separating sections.

### Immunohistochemistry (IHC)

*PPG*^*YFP*^ NTS tissue was processed free-floating and incubated with chicken anti-green fluorescent protein (*GFP*; AbCam, ab13970, 1:1000) and/or rabbit anti-c-FOS (Calbiochem, PC38, 1:5000) primary antibodies. This anti-*GFP* antibody binds selectively to YFP in this mouse (21). YFP was visualized fluorometrically with an anti-chicken Alexa Fluor 488 (1:500, Jackson ImmunoResearch Laboratories) secondary antibody (1:500, Life Technologies) and c-FOS chromogenically with a biotinylated donkey anti-rabbit (1:500, Jackson ImmunoResearch Laboratories) and diaminobenzidine (DAB) as previously described (38, 39). For the c-FOS study, 10 mice were injected i.p. with either saline or 10 mg/kg lorcaserin and perfuse-fixed with 4% paraformaldehyde (PFA) 2 hours later. Brains were excised and post-fixed overnight in 4% PFA at 6°C. Brains were stored in PBS until IHC processing.

### In Situ Hybridization (ISH) and IHC

Fluorescent ISH (FISH) using digoxigenin (DIG)-UTP labelled RNA probes was performed in conjunction with IHC. Specifically, to label *Htr2c* (5-HT_2C_R) mRNA-expressing neurons, a linearized expression vector (pBluescript SK-) containing 3 kb of rat 5-HT_2C_R cDNA (XM_01761920.1: 28-3006) was used as DNA template for in vitro transcription (40). To generate a DNA template for labelling *Ppg* mRNA-expressing neurons, PPG cDNA (NM_008100.2: 136 to 946) was PCR-amplified from mouse brainstem cDNA. *In vitro* transcription of antisense and sense probes was performed with a DIG-RNA Labelling Mix (Roche, Mannheim, Germany) using either T7 or T3 polymerase.

FISH and IHC in the same tissue were performed with either IHC (free floating) then FISH (NTS sections mounted on slides) or FISH then IHC. In POMC mice, red fluorescent protein (RFP) IHC was performed followed by *Ppg*^*dsRed*^ FISH. In *MC4R*^*GFP*^ mice, *GFP* IHC was conducted and then *Ppg* FISH. In *Ppg*^*YFP*^ mice, *Ppg* FISH was performed followed by *GFP* IHC and 5-ht2Cr FISH was conducted followed by *GFP* IHC. In *5-HT2CR*^*Cre:YFP*^ mice ARC-injected with anterograde tracer AAV9.CAG.Flex.tdTomato.WPRE.bGH, PPG FISH was performed followed by RFP IHC. NTS sections on slides were incubated with 1% sodium borohydride (15 min), 0.1 M Triethanolamine (TEA) (20 min), 0.25% Acetic Anhydride in 0.1 M TEA (10 min) and washed in saline sodium citrate (SSC) buffer (41). Sections were then preincubated in hybridization buffer (1 h) followed by overnight incubation with DIG-UTP-labelled probes (1 or 2 ng/μl) in hybridization buffer at 72°C. The next day, sections were washed with 2xSSC and incubated in 20 μg/ml RNase A (30 min), 1xSSC (15 min), 0.1xSSC (30 min), 1% H_2_O_2_ (15 min), wash buffer (100 mM Tris, pH 7.5, 150 mM NaCl; 3x 5 min) and immersed in a blocking buffer (100 mM Tris, pH 7.5, 150 mM NaCl, 0.5% blocking reagent FP1020, Perkin-Elmer, Boston, USA; 1 h). Sections were then incubated overnight with a peroxidase conjugated anti-DIG antibody (1:200, Roche, Mannheim, Germany) and primary antibodies for *GFP*/YFP (chicken anti-*GFP*, 1:1000, ab13970, AbCam, Cambridge UK) or dsRed/tdTomato (Rabbit anti-RFP, 1:500, 600-401-379, Rockland Immunochemicals, Inc, Limerick PA, USA). The following day, the sections were treated with TSA PLUS Biotin Kit (PerkinElmer Inc., Waltham, MA USA) and the respective mRNAs (*5-ht2Cr* or *Ppg*) visualized with streptavidin conjugated Alexa Fluor 488 (1:500, S11223, Life Technologies, Carlsbad, CA USA) or 568 (1:500, 1024067, Life Technologies). IHC was performed concomitantly with the FISH detection with anti-rabbit Alexa Fluor 568 (1:500, A10042, Life Technologies, Carlsbad, CA USA) or anti-chicken Alexa Fluor 488 (1:500, 703-545-155, Jackson ImmunoResearch Laboratories) secondary antibodies.

Single label ISH for *Glp1r* mRNA was performed with DAB to quantify GLP-1R knockdown in mice treated with AAV1-shGlp1r-*GFP* or control AAV1-shRNA-*GFP*. Single label ISH for *Glp1r* mRNA was performed as above except for overnight hybridization incubation which was performed at 62°C. Following TSA PLUS Biotin Kit treatment, sections were incubated in wash buffer (100 mM Tris, pH 7.5, 150 mM NaCl; 3x 5 min) followed by incubation in streptavidin-HRP (1:250, FP1047, Perkin-Elmer, Boston, USA; 1 h). Sections were then treated with Vectastain® Elite ABC Kit (PK-6100, Vector Laboratories Inc., California, USA) and visualized using ImmPACT® DAB EqV Substrate Kit (SK-4103, Vector Laboratories Inc., California, USA).

Dual ISH and IHC in PPG-YFP tissue was performed with a ^35^S UTP radio-labelled PPG probe to detect PPG mRNA and a goat α-*GFP* antibody (Abcam, 1:1000) to visualize YFP as previously described (14, 41).

### 3D rendering of NTS IHC and FISH signal expression

3D models for the distribution of PPG^YFP^-IR + *5-ht*_*2C*_ *r* mRNA (FISH), POMC^dsRed^-IR + *Ppg* mRNA (FISH) and MC4R^*GFP*^-IR + *Ppg* mRNA (FISH) were generated by assembling 11 to 14 consecutive NTS sections spaced 125 μm apart with Free D modelling software (42). The NTS, the surface of the brain and the central canal were outlined for each section, and the anatomical location of individual FISH or IHC labelled cells was entered manually. Colours and cell sizes were assigned in close resemblance to the structures in the digitized images, with single-labelled cells shown in red and green and dual-labelled cells shown in yellow. 3D models are presented in a caudal to rostral orientation.

### Viral vectors

For the ablation of PPG neurones in the NTS we used AAV8-FLEX-mCherry-DTA and AAV8-FLEX-E*GFP* (UNC, ETHZ; Table 2). For the chemogenetic activation of 5-HT2cR we used AAV8-hSyn-DIO-hMD3q-mCherry and AAV8-hSyn-DIO-mCherry (UNC, Addgene). For the silencing of GLP1R mRNA we used AAV1.U6.shRGlp1r07.CB7.E*GFP*.SV40 (AAV1-shRNA-Glp1r) and control AAV1.CB7.Cl.e*GFP*.WPRE.rBG (AAV1-shRNA-Control) (Perelman School of Medicine, University of Pennsylvania). For the ARC neuronal projections study, AAV9.CAG.flex.tdTomato.WPRE.bGH (Perelman School of Medicine, University of Pennsylvania; Virus titer 1.55e13 GC/ml) was used.

**Table 2.**
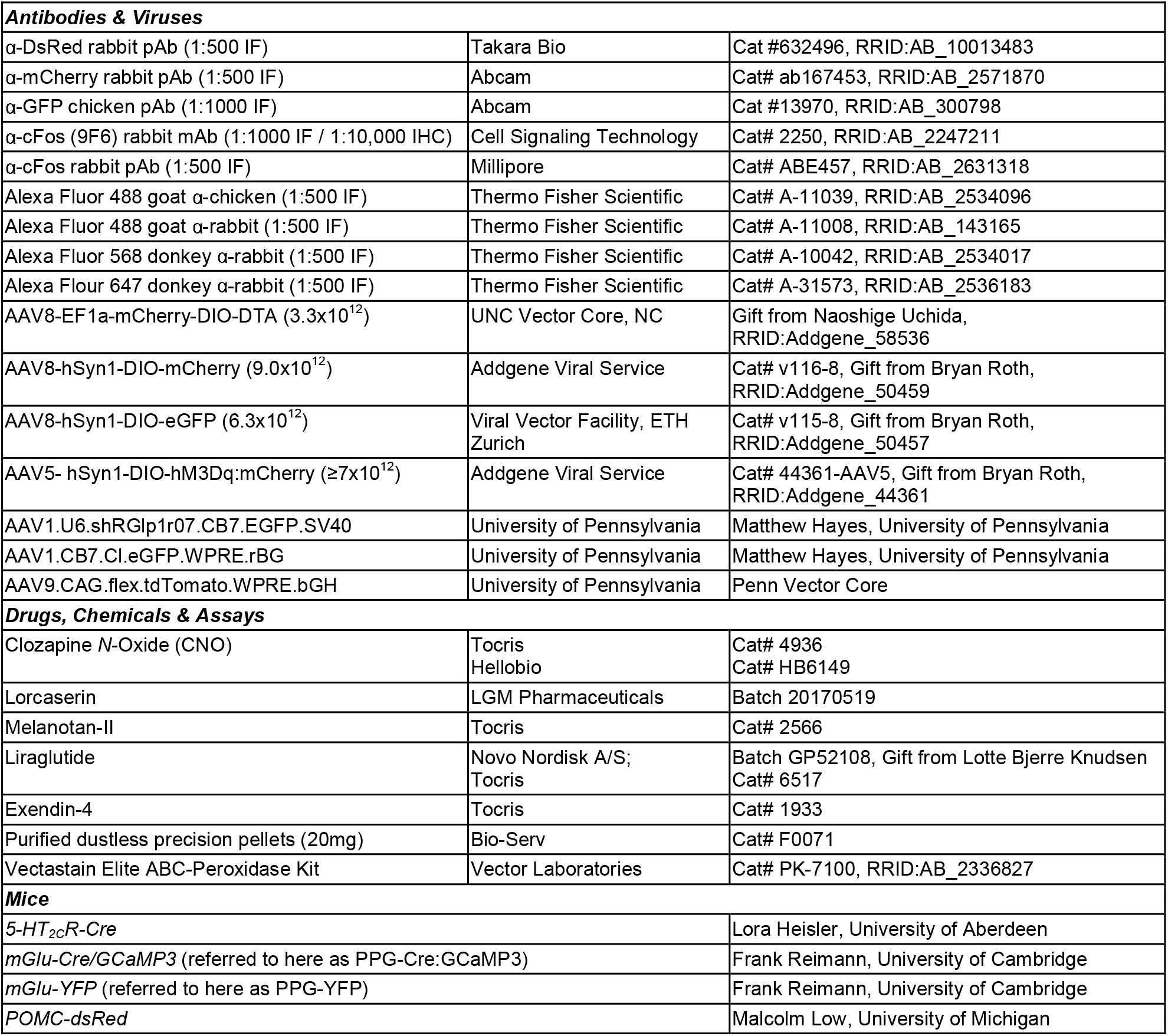
Reagents and resources

### Ablation of PPG^NTS^ neurons

*Ppg*^*CRE*^ mice were anaesthetised with i.m. ketamine hydrochloride (50 mg/kg) and medetomidine (1 mg/kg) with peri-operative carprofen analgesia (5 mg/kg s.c.). The skull was fixed in a stereotaxic frame and the head ventroflexed to create a right angle between the nose and the neck. Neck muscles were bisected along the midline from the occipital crest to the first vertebrae, and the atlanto-occipital membrane was horizontal bisected to reveal the dorsal brainstem surface and obex. Using a glass microcapillary pulled to a 40 μm tip diameter, 250 nl of virus (AAV8-mCherry-DIO-DTA or AAV8-DIO- E*GFP*) was injected slowly (50 nl/min) bilaterally at the following coordinates relative to obex: 0.50 mm lateral, 0.10 mm rostral, and 0.35 mm ventral. At the end of experiments expression of viral transgenes and successful ablation was confirmed immunohistologically for all experimental animals included in analyses as described previously (11). PPG cell bodies were absent from the caudal NTS of AAV8-mCherry-DIO-DTA injected mice, whilst PPG cells in the IRt remained intact in these animals (Figure 2k; Supplementary Figure 2d-j).

### Chemogenetics, mRNA silencing and neuronal projections studies

For DREADD, shGlp1r (generous gift of Matthew Hayes, University of Pennsylvania) and retrograde tracer studies, methods were used as previously described (5, 43, 44). Briefly, adult male *5-HT2CR*^*CRE*^ mice were anaesthetised with Isoflurane (5% induction, 2% maintenance) with perioperative analgesia (carprofen 5mg/Kg, SC) before and during 48h after the surgery and given at least 14 days of recovery before any treatment. For the delivery of AAVs into the NTS, mice were injected bilaterally as described in the section above using a pneumatic injector (Harvard Apparatus, Cambridge, UK). Tissue was processed with IHC for fluorescent proteins mCherry or YFP to visualise and confirm veracity of injection sites (e.g. Figure 2j-k, 3a). Mice with injections outside of the NTS were excluded from analysis. ISH with DAB as the chromogen for *Glp1r* mRNA was performed using methods described above on tissue from mice injected with AAV1-shRNA-control or AAV1-shRNA-Glp1r. Estimates of NTS and AP GLP-1R knockdown was quantified using densitometry analysis (NIH ImageJ). To visualise neuronal projections from the ARC to the NTS, mice were injected unilaterally in coordinates -1.58 mm from bregma, 5.75 mm depth from brain surface and .2 mm from midline. Tissue was processed for dsRed-IR and *Ppg* FISH, as detailed above in ISH and IHC section.

### Feeding studies

The feeding behaviour experiments in PPG^NTS^ ablated mice were conducted using a mixed model design. Mice in control (e*GFP*-transduced) and ablated (DTA-transduced) cohorts each received both saline and test drugs (i.e. lorcaserin and MTII respectively) in a counterbalanced order separated by a minimum 72 hour washout period. To prevent any confound from stress-induced hypophagia, mice were extensively habituated to single-housing, handling, saline dosing and the food measurement protocol, such that baseline intakes were stable for ≥ 3 saline-dosed habituation sessions, prior to commencement of test drug sessions. On habituation and test days, mice were weighed and food removed for 3 hours prior to dark onset, such that they were entrained to eat consistently at dark onset, but without inducing a negative energy balance that would elicit a large refeed response to physiologically recruit PPG^NTS^ neurons (11). Mice were administered obesity medications or saline vehicle i.p. at 5 mg/ml dose volume 30 minutes prior to dark onset, at which point a pre-weighed amount of their standard chow (Teklad 2018, Envigo) was returned, and intake measured 1, 2, 4 and 21 hours later, at which point 24 hr body weight change was also measured.

All other feeding studies were performed using a similar design, with intake assessed up to 6 hours, unless otherwise specified. A subset of MC4R wild type control littermates were fed 60%-fat diet (HFD; Test Diet, 58Y1) from 3 weeks of age to match *MC4R* knockout obesity. All other genotypes were fed chow throughout.

### Drugs

Drugs were prepared in sterile saline. Lorcaserin (LGM Pharmaceuticals) was administered at 2.5 to 10 mg/kg, i.p.; Exendin-4 (Tocris) was administered at 0.004 to 0.04 mg/kg, i.p.; Liraglutide (Tocris and Novo Nordisk) was administered at 0.01 to 0.2 mg/kg, s.c or i.p. and MTII (Tocris) was injected at 3 mg/kg, i.p.; Clozapine N-oxide (CNO) (Tocris and Hellobio) was administered at 1mg/kg, i.p..

### Statistical analyses

Raw data were processed in Microsoft Excel, plots were generated in GraphPad Prism 7.0 and statistical analyses conducted in SPSS 26.0 (IBM Corp, NY, USA). For transparency, food intake data are presented as individual data points, including within-subjects relationships where appropriate, and summary data are presented as mean ± SEM. Food intake data were initially analysed by two- or three-way mixed model ANOVA, with drug treatment as the within-subjects factor and virus (or genotype) and sex as between-subjects factors. No significant interactions with sex were observed. Data from all mice were analysed with treatment x virus (or genotype) mixed model two-way ANOVA or one-way within-subjects ANOVA for lorcaserin dose-response and loracserin + liraglutide combination study. Where sphericity could not be assumed for repeated-measures ANOVA, the Geisser-Greenhouse adjustment was applied, in which case the revised fractional degrees of freedom are reported in figure legends. Where treatment x virus interactions were significant (at p<0.05), simple main effects with Sidak’s correction for multiple comparisons were performed. Where no significant interaction was observed, main effects of treatment were presented. Cell counts for FOS-IR were analysed by unpaired Student’s t-test (with Welch’s correction as appropriate) or Mann-Whitney test. Statistics are presented in Figure legends and data presented as mean ± SEM.

### Ethical Approval

All experiments were performed in accordance with the U.K. Animals (Scientific Procedures) Act, 1986 and received local and Home Office ethical approval.

## FIGURE LEGENDS

**Supplementary Figure 1.**
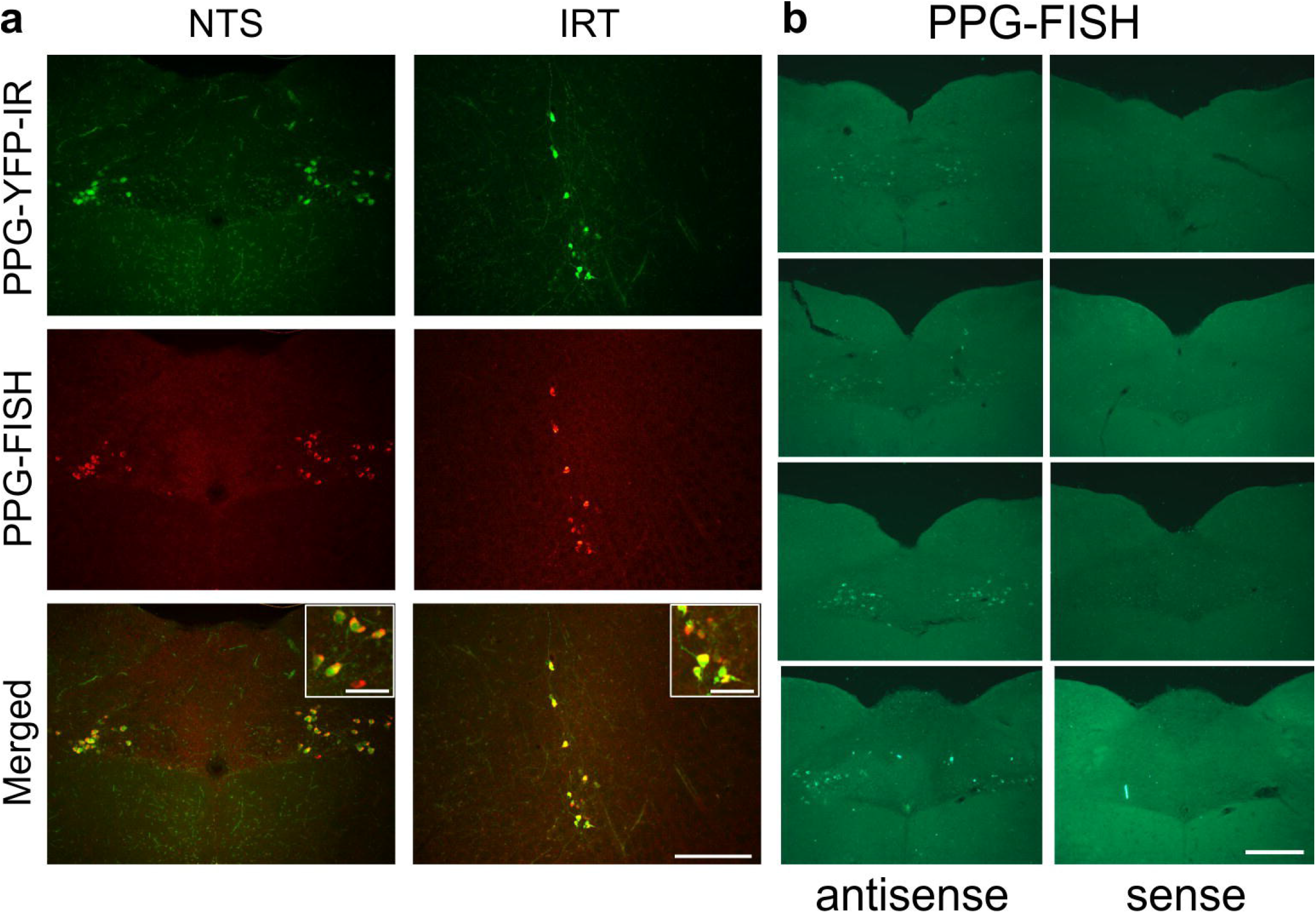
PPG expression within the NTS. (a) Representative photomicrographs illustrating that pre-proglucagon yellow fluorescent protein immunoreactivity (PPG-YFP-IR; green, top panels) and *Ppg* mRNA using fluorescent in situ hybridization (FISH; antisense probe targeting *Ppg* mRNA NM_008100.2: nucleotides 136 to 946; red, middle panels) neurons colocalize (yellow, bottom panels) in the nucleus of the solitary tract (NTS; upper panels) and intermediate reticular nucleus (IRt; lower panels) in *Ppg*^*YFP*^ mice (n=4). CC, central canal; scale bars, 200 μm and 50 μm (insets), respectively. (b) Representative photomicrographs of *Ppg* FISH using an antisense probe (targeting *Ppg* mRNA NM_008100.2: nucleotides 136 to 946; left panels) or control sense probe (right panels) in the caudal to rostral NTS (−7.8 to -7.5 from bregma) of 5-HT R_2C_^CRE:YFP^ mice (n=4). *Ppg* FISH antisense, but not control sense, labels PPG neurons in green. Intrinsic 5-HT R_2C_^CRE:YFP^ fluorescence is not detected following perfusion fixation and FISH processing in NTS tissue. CC – central canal. Scale bar: 200 μm.

**Supplementary Figure 2.**
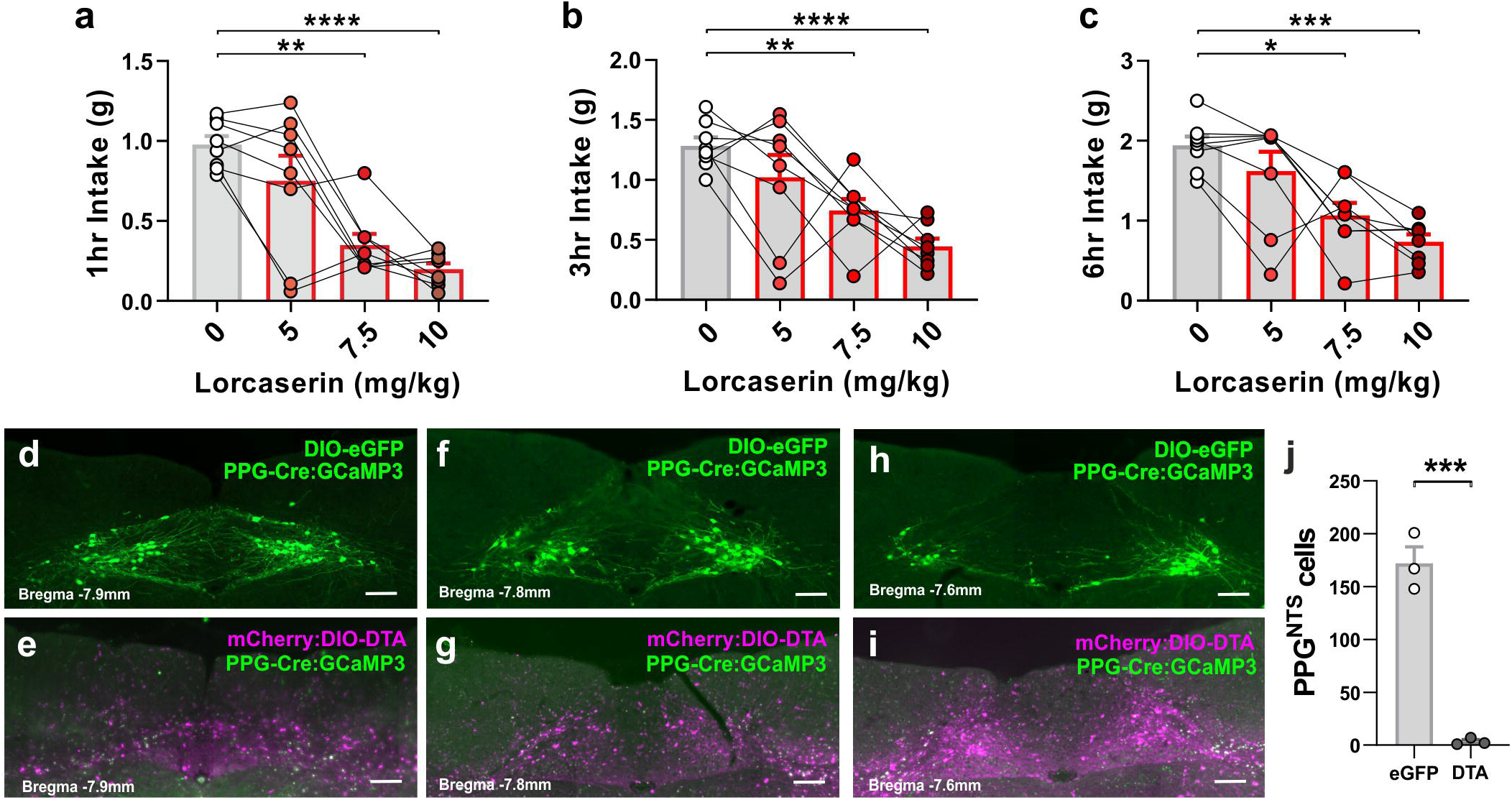
Dose response of lorcaserin and DTA ablation of PPG^NTS^ neurons. (a-c) Dose response of lorcaserin (5, 7.5, or 10 mg/kg i.p.) on dark onset food intake in male and female control mice (n=8). Lorcaserin dose-dependently decreased food intake at 1, 3 and 6 hours (1hr: treatment F_(1.6, 11.1)_=18.54, p=0.0005; 3hr: F_(1.8, 12.3)_=10.36, p=0.0029; 6hr: F_(1.4, 10.1)_=11.47, p=0.0041). Data are presented as mean ± SEM; ** p < 0.01, *** p < 0.001, **** p<0.0001. (d-j) The nucleus of the solitary tract (NTS) of PPG-Cre:GCaMP3 mice was injected bilaterally with either AAV8-mCherry:DIO-DTA to ablate PPG neurons or with AAV8-DIO-eGFP as control. (d-i) Representative photomicrographs at three rostrocaudal levels showing the presence or absence of PPG^NTS^ neurons, respectively, after transduction with control or DTA virus. Magenta shows spread of the DTA virus by visualising the Cre-independent transduction of mCherry. (j) Quantification of PPG^NTS^ neurons in randomly selected eGFP and DTA transduced mice (n=3), respectively. Scale bars in d-i: 100 μm.

**Supplementary Figure 3.**
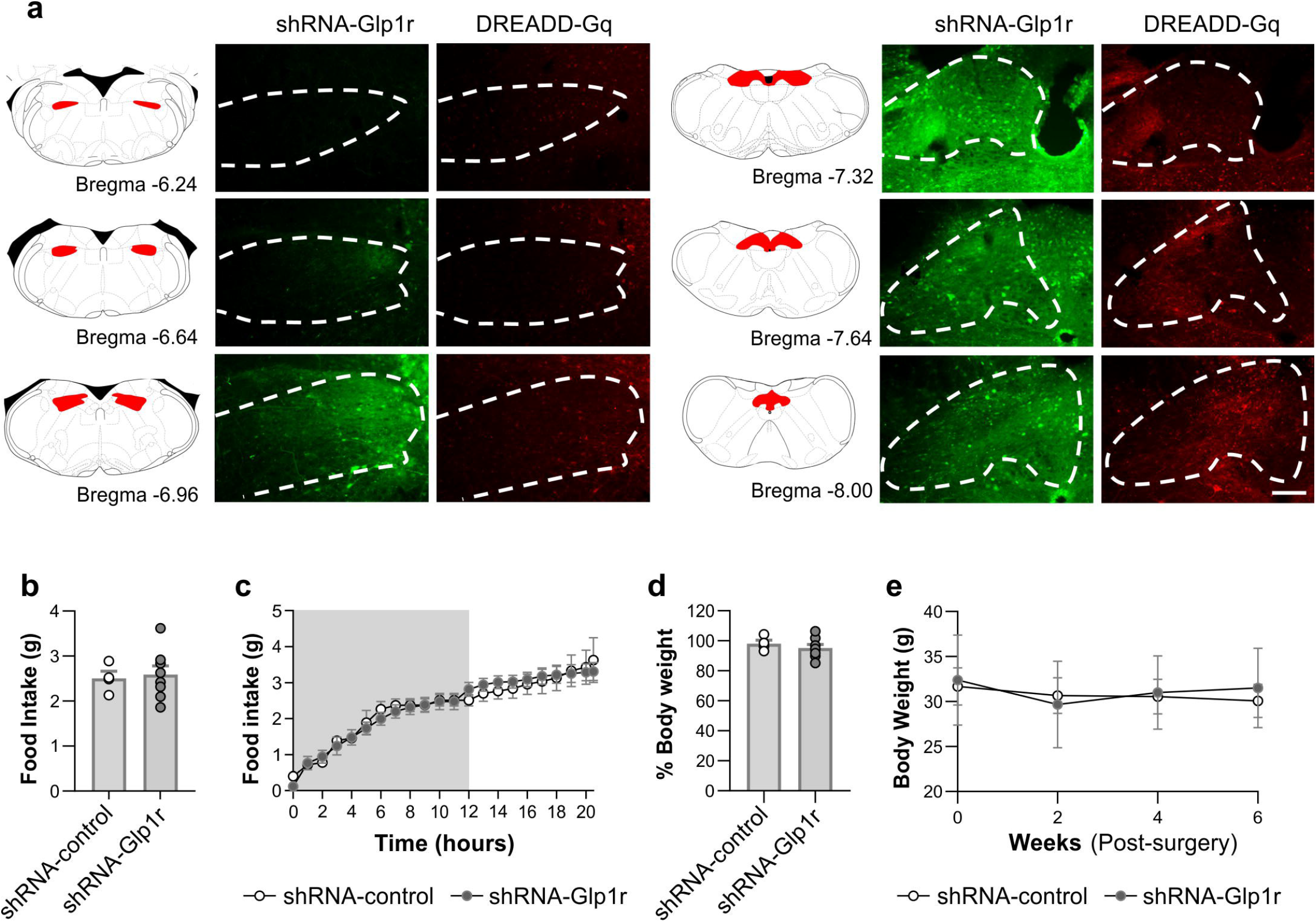
Glp-1r knockdown in the NTS does not alter food intake or body weight. *5-HT*_*2C*_*R*^*Cre*^ mice received NTS injections of a viral cocktail comprising AAV8-DIO-hM3Dq-mCherry (or DIO-mCherry as a control) to evaluate the effect of 5-HT_2C_R^NTS^ neuronal activation, and AAV1-shGlp1r-*GFP* (or AAV1-*GFP* as a control) to assess the importance of GLP-1R^NTS^ in feeding effects induced by chemogenetic activation of 5-HT R_2C_^NTS^ neurons. (a) Representative images of NTS injection localization and spread for mCherry and *GFP* from the viruses infused. Scale bar = 100μm. Mice were tracked for 6 weeks post-treatment. Knockdown of GLP-1R within the NTS had no effect on (b-c) *ad libitum* food intake or (d-e) body weight. Data presented as mean ± SEM.

**Supplementary Figure 4.**
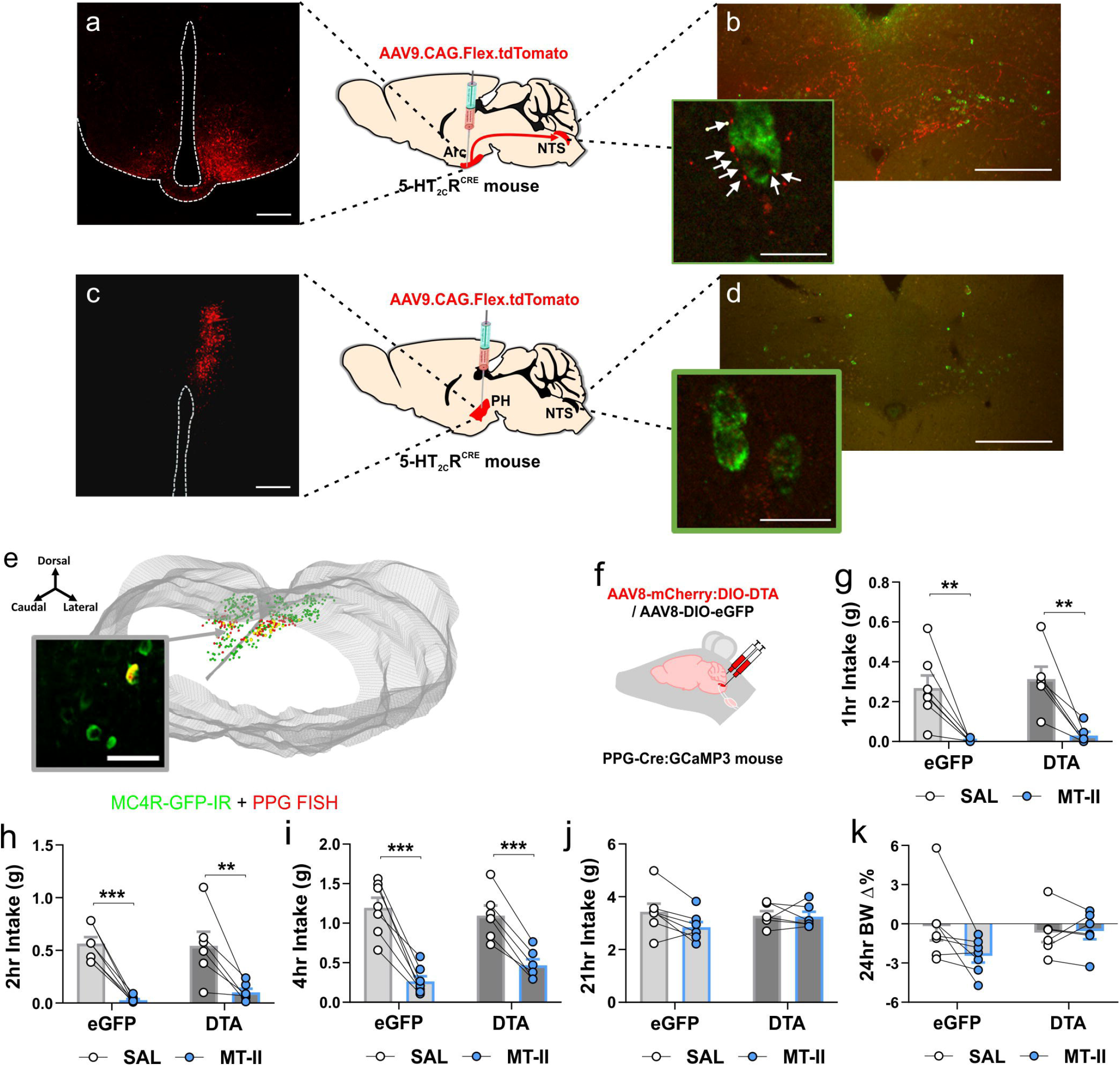
Arcuate nucleus of the hypothalamus 5-HT_2C_R neurons project to PPG^NTS^ neurons and PPG^NTS^ ablation does not affect MC4R agonist hypophagia. (a-d) AAV9.CAG.Flex.tdTomato.WPRE.bGH (red) injected into hypothalamic subregions of 5-HT_2C_R mice (n=10) to visualize axon terminals within the brain. (a) Arcuate nucleus of the hypothalamus (Arc) injections (n=4 mice) revealed (b) dense innervation of the nucleus of the solitary tract (NTS) that was not seen following injections outside the Arc (n=6 mice). Arc derived *5-HT*_*2C*_*R*^*Cre:YPF*^ terminals (red) were in close apposition to a subset of *Ppg* mRNA neurons (green) vizualized with fluorescent in situ hybridization histochemistry (FISH). (c) Representative photomicrograph of AAV9.CAG.Flex.tdTomato.WPRE.bGH injected into the posterior hypothalamus (PH) and (d) corresponding limited innervation of the NTS (PPG^NTS^ neurons, green). (e) 3D rendering of the caudal to rostral NTS from *MC4R*^*GFP*^ mice processed for FISH for *Ppg* mRNA (red cells) and *GFP*-IR (green cells) illustrating limited overlap (13% of PPG^NTS^ co-express MC4Rs, n=4 mice). 3V, third ventricle; scale bars a-d, 200 μm inset, 25 μm. (f-k) Dark onset food intakes and body weight changes in male and female PPG^NTS^ DTA-ablated (n=6) and eGFP (n=7) control *Ppg*^Cre^ mice administered saline or preclinical obesity medication MC4R agonist melanotan-II (MTII; 3 mg/kg i.p.). (f) Schematic showing the injection site of viral delivery into the NTS. MTII reduced cumulative chow intake in both PPG^NTS^ neuron ablated and control mice at (g) 1 hour (treatment F_(1, 11)_=38.47, p<0.0001; treatment x virus F_(1,11)_=0.04641, p=0.8334), (h) 2 hours (treatment F_(1, 11)_=57.07, p<0.0001; treatment x virus F_(1,11)_ = 0.5720, p=0.4654) and (i) 4 hours (treatment F_(1, 11)_=108.8, p<0.0001; treatment x virus F_(1,11)_=4.105, p=0.0677). However, MTII did not significantly reduce intake in either group at (j) 21 hours or have an effect on (k) 24 hour body weight change. Data are presented as mean±SEM; * p < 0.05, *** p < 0.001.

**Supplementary Figure 5.**
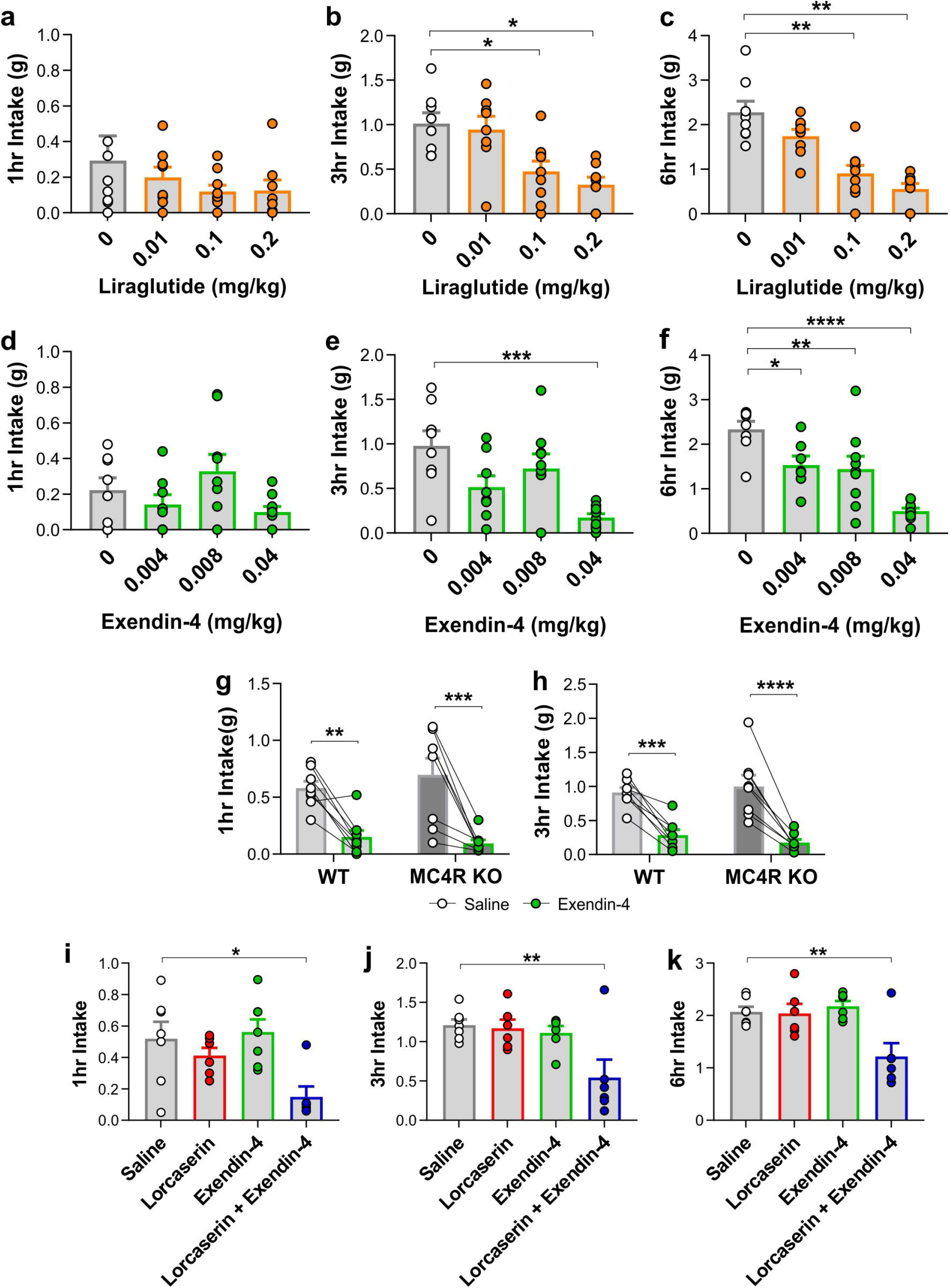
Combination of lorcaserin and exendin-4 significantly reduce food intake. (a-c) Dose response of liraglutide on dark onset food intake in male and female control mice (n=8) administered saline or liraglutide (0.01, 0.1, or 0.2 mg/kg s.c.). Liraglutide significantly reduced cumulative food intake at 0.1mg/kg and 0.2mg/kg at (b) 3 hours (treatment F_(3,29)_=8.040, p=0.0005), and (c) 6 hours (treatment F_(3,29)_=17.71, p<0.0001). (d-f) Dose response of exendin-4 on food intake in male and female control mice (n=8) administered saline or exendin-4 (0.004, 0.008, or 0.04 mg/kg i.p.). Exendin-4 significantly reduced cumulative food intake at 0.008mg/kg and 0.04mg/kg at (e) 3 hours (treatment, F_(3,27)_=9.672, p=0.0002), and (f) 6 hours (treatment F_(3,29)_=21.34, p<0.0001). Data were analysed by one-way ANOVA. Significant treatment interactions were followed by analysis Dunnett’s multiple comparison test. Exendin-4 (0.04 mg/kg, i.p.) signficantly reduced (g) 1 and (h) 3 hour cumulative dark cycle food intake in both male and female wild type and *Mc4r* knockout mice. (n=8; 1hr: treatment F_(1,14)_=41.71, p<0.0001, treatment x genotype F_(1,14)_=1.164, p=0.299; 3hr: treatment F_(1,14)_=84.12, p<0.0001,treatment x genotype F_(1,14)_=1.571, p=0.231,). (i-k) Mice were fasted during the dark cycle followed by I.P injection of saline (n=7), lorcaserin (5 mg/kg, n=8), exendin-4 (0.004 mg/kg, n=6) or a combination of lorcaserin (5 mg/kg) and exendin-4 (0.004 mg/kg) (n=6). Food intake was measured at 1, 3, and 6 hours. No significant reduction in food intake was measured following treatment with lorcaserin (5 mg/kg) or exendin-4 (0.004 mg/kg) alone at any time point. A significant reduction in food intake following combination treatment was observed at 1 hour (treatment F_(3,22)_=4.942, p=0.009), 3 hours (treatment F_(3,21)_=5.288, P=0.007), and 6 hours (treatment F_(3,21)_=6.877, P=0.002). All data are presented as mean ± SEM; * p<0.05, ** p<0.01, *** p<0.001, **** p<0.0001.

## REFERENCES

1. Secher A, Jelsing J, Baquero AF, Hecksher-Sorensen J, Cowley MA, Dalboge LS, et al. The arcuate nucleus mediates GLP-1 receptor agonist liraglutide-dependent weight loss. J Clin Invest. 2014;124(10):4473–88.

2. Shah BP, Vong L, Olson DP, Koda S, Krashes MJ, Ye C, et al. MC4R-expressing glutamatergic neurons in the paraventricular hypothalamus regulate feeding and are synaptically connected to the parabrachial nucleus. Proc Natl Acad Sci U S A. 2014;111(36):13193–8.

3. Balthasar N, Dalgaard LT, Lee CE, Yu J, Funahashi H, Williams T, et al. Divergence of melanocortin pathways in the control of food intake and energy expenditure. Cell. 2005;123(3):493–505.

4. Li MM, Madara JC, Steger JS, Krashes MJ, Balthasar N, Campbell JN, et al. The Paraventricular Hypothalamus Regulates Satiety and Prevents Obesity via Two Genetically Distinct Circuits. Neuron. 2019;102(3):653–67 e6.

5. D’Agostino G, Lyons D, Cristiano C, Lettieri M, Olarte-Sanchez C, Burke LK, et al. Nucleus of the Solitary Tract Serotonin 5-HT2C Receptors Modulate Food Intake. Cell Metab. 2018;28(4):619–30 e5.

6. Burke LK, Ogunnowo-Bada E, Georgescu T, Cristiano C, de Morentin Pbm, Valencia Torres L, et al. Lorcaserin improves glycemic control via a melanocortin neurocircuit. Molecular metabolism. 2017;6(10):1092–102.

7. Leon RM, Borner T, Reiner DJ, Stein LM, Lhamo R, De Jonghe BC, et al. Hypophagia induced by hindbrain serotonin is mediated through central GLP-1 signaling and involves 5-HT2C and 5-HT3 receptor activation. Neuropsychopharmacology : official publication of the American College of Neuropsychopharmacology. 2019.

8. Stemmer K, Muller TD, DiMarchi RD, Pfluger PT, and Tschop MH. CNS-targeting pharmacological interventions for the metabolic syndrome. J Clin Invest. 2019;130:4058–71.

9. Gaykema RP, Newmyer BA, Ottolini M, Raje V, Warthen DM, Lambeth PS, et al. Activation of murine pre-proglucagon-producing neurons reduces food intake and body weight. J Clin Invest. 2017;127:1031–45.

10. Shi X, Chacko S, Li F, Li D, Burrin D, Chan L, et al. Acute activation of GLP-1-expressing neurons promotes glucose homeostasis and insulin sensitivity. Molecular metabolism. 2017;6(11):1350–9.

11. Holt MK, Richards JE, Cook DR, Brierley DI, Williams DL, Reimann F, et al. Preproglucagon Neurons in the Nucleus of the Solitary Tract Are the Main Source of Brain GLP-1, Mediate Stress-Induced Hypophagia, and Limit Unusually Large Intakes of Food. Diabetes. 2019;68(1):21–33.

12. Brierley DI, Holt MK, Singh A, de Araujo A, McDougle M, Vergara M, et al. Central and peripheral GLP-1 systems independently suppress eating. Nat Metab. 2021;3(2):258–73.

13. Hisadome K, Reimann F, Gribble FM, and Trapp S. Leptin directly depolarizes preproglucagon neurons in the nucleus tractus solitarius: electrical properties of glucagon-like Peptide 1 neurons. Diabetes. 2010;59(8):1890–8.

14. Garfield AS, Patterson C, Skora S, Gribble FM, Reimann F, Evans ML, et al. Neurochemical characterization of body weight-regulating leptin receptor neurons in the nucleus of the solitary tract. Endocrinology. 2012;153(10):4600–7.

15. Holt MK, Llewellyn-Smith IJ, Reimann F, Gribble FM, and Trapp S. Serotonergic modulation of the activity of GLP-1 producing neurons in the nucleus of the solitary tract in mouse. Molecular metabolism. 2017;6(8):909–21.

16. Sharma S, Garfield AS, Shah B, Kleyn P, Ichetovkin I, Moeller IH, et al. Current Mechanistic and Pharmacodynamic Understanding of Melanocortin-4 Receptor Activation. Molecules. 2019;24(10).

17. Marston OJ, Garfield AS, and Heisler LK. Role of central serotonin and melanocortin systems in the control of energy balance. Eur J Pharmacol. 2011;660(1):70–9.

18. Dowsett GKC, Lam BYH, Tadross JA, Cimino I, Rimmington D, Coll AP, et al. A survey of the mouse hindbrain in the fed and fasted state using single-nucleus RNA sequencing. Molecular metabolism. 2021:101240.

19. Zheng H, Stornetta RL, Agassandian K, and Rinaman L. Glutamatergic phenotype of glucagon-like peptide 1 neurons in the caudal nucleus of the solitary tract in rats. Brain structure & function. 2014.

20. Trapp S, and Cork SC. PPG neurons of the lower brain stem and their role in brain GLP-1 receptor activation. Am J Physiol Regul Integr Comp Physiol. 2015;309(8):R795–804.

21. Llewellyn-Smith IJ, Reimann F, Gribble FM, and Trapp S. Preproglucagon neurons project widely to autonomic control areas in the mouse brain. Neuroscience. 2011;180:111–21.

22. Xu Y, Jones JE, Kohno D, Williams KW, Lee CE, Choi MJ, et al. 5-HT2CRs expressed by pro-opiomelanocortin neurons regulate energy homeostasis. Neuron. 2008;60(4):582–9.

23. Berglund ED, Liu C, Sohn JW, Liu T, Kim MH, Lee CE, et al. Serotonin 2C receptors in pro-opiomelanocortin neurons regulate energy and glucose homeostasis. J Clin Invest. 2013;123(12):5061–70.

24. Heisler LK, and Lam DD. An appetite for life: brain regulation of hunger and satiety. Curr Opin Pharmacol. 2017;37:100–6.

25. Ghamari-Langroudi M, Digby GJ, Sebag JA, Millhauser GL, Palomino R, Matthews R, et al. G-protein-independent coupling of MC4R to Kir7.1 in hypothalamic neurons. Nature. 2015;520(7545):94–8.

26. Hisadome K, Reimann F, Gribble FM, and Trapp S. CCK Stimulation of GLP-1 Neurons Involves {alpha}1-Adrenoceptor-Mediated Increase in Glutamatergic Synaptic Inputs. Diabetes. 2011;60(11):2701–9.

27. Iepsen EW, Zhang J, Thomsen HS, Hansen EL, Hollensted M, Madsbad S, et al. Patients with Obesity Caused by Melanocortin-4 Receptor Mutations Can Be Treated with a Glucagon-like Peptide-1 Receptor Agonist. Cell Metab. 2018;28(1):23–32 e3.

28. Adams JM, Pei H, Sandoval DA, Seeley RJ, Chang RB, Liberles SD, et al. Liraglutide Modulates Appetite and Body Weight Through Glucagon-Like Peptide 1 Receptor-Expressing Glutamatergic Neurons. Diabetes. 2018;67(8):1538–48.

29. Fortin SM, Lipsky RK, Lhamo R, Chen J, Kim E, Borner T, et al. GABA neurons in the nucleus tractus solitarius express GLP-1 receptors and mediate anorectic effects of liraglutide in rats. Sci Transl Med. 2020;12(533).

30. van Can J, Sloth B, Jensen CB, Flint A, Blaak EE, and Saris WH. Effects of the once-daily GLP-1 analog liraglutide on gastric emptying, glycemic parameters, appetite and energy metabolism in obese, non-diabetic adults. Int J Obes (Lond). 2014;38(6):784–93.

31. Reimann F, Habib AM, Tolhurst G, Parker HE, Rogers GJ, and Gribble FM. Glucose sensing in L cells: a primary cell study. Cell Metab. 2008;8(6):532–9.

32. Parker HE, Adriaenssens A, Rogers G, Richards P, Koepsell H, Reimann F, et al. Predominant role of active versus facilitative glucose transport for glucagon-like peptide-1 secretion. Diabetologia. 2012;55(9):2445–55.

33. Burke LK, Doslikova B, D’Agostino G, Greenwald-Yarnell M, Georgescu T, Chianese R, et al. Sex difference in physical activity, energy expenditure and obesity driven by a subpopulation of hypothalamic POMC neurons. Molecular metabolism. 2016;5(3):245–52.

34. Hentges ST, Otero-Corchon V, Pennock RL, King CM, and Low MJ. Proopiomelanocortin expression in both GABA and glutamate neurons. J Neurosci. 2009;29(43):13684–90.

35. Stuart T, Butler A, Hoffman P, Hafemeister C, Papalexi E, Mauck WM, 3rd, et al. Comprehensive Integration of Single-Cell Data. Cell. 2019;177(7):1888–902 e21.

36. Butler A, Hoffman P, Smibert P, Papalexi E, and Satija R. Integrating single-cell transcriptomic data across different conditions, technologies, and species. Nat Biotechnol. 2018;36(5):411–20.

37. Burke LK, Doslikova B, D’Agostino G, Garfield AS, Farooq G, Burdakov D, et al. 5-HT obesity medication efficacy via POMC activation is maintained during aging. Endocrinology. 2014;155(10):3732–8.

38. Lam DD, Zhou L, Vegge A, Xiu PY, Christensen BT, Osundiji MA, et al. Distribution and neurochemical characterization of neurons within the nucleus of the solitary tract responsive to serotonin agonist-induced hypophagia. Behav Brain Res. 2009;196(1):139–43.

39. Heisler LK, Jobst EE, Sutton GM, Zhou L, Borok E, Thornton-Jones Z, et al. Serotonin reciprocally regulates melanocortin neurons to modulate food intake. Neuron. 2006;51(2):239–49.

40. Julius D, MacDermott AB, Axel R, and Jessell TM. Molecular characterization of a functional cDNA encoding the serotonin 1c receptor. Science. 1988;241(4865):558–64.

41. Elias CF, Aschkenasi C, Lee C, Kelly J, Ahima RS, Bjorbaek C, et al. Leptin differentially regulates NPY and POMC neurons projecting to the lateral hypothalamic area. Neuron. 1999;23(4):775–86.

42. Schwarz J, Burguet J, Rampin O, Fromentin G, Andrey P, Tome D, et al. Three-dimensional macronutrient-associated Fos expression patterns in the mouse brainstem. PLoS One. 2010;5(2):e8974.

43. D’Agostino G, Lyons DJ, Cristiano C, Burke LK, Madara JC, Campbell JN, et al. Appetite controlled by a cholecystokinin nucleus of the solitary tract to hypothalamus neurocircuit. Elife. 2016;5.

44. Schmidt HD, Mietlicki-Baase EG, Ige KY, Maurer JJ, Reiner DJ, Zimmer DJ, et al. Glucagon-Like Peptide-1 Receptor Activation in the Ventral Tegmental Area Decreases the Reinforcing Efficacy of Cocaine. Neuropsychopharmacology : official publication of the American College of Neuropsychopharmacology. 2016;41(7):1917–28.

